# Maternal transmission of a plastid structure enhances offspring fitness

**DOI:** 10.1101/2025.10.21.683706

**Authors:** Tyler J. Carrier, Andrés Rufino-Navarro, Thorben Knoop, Urska Repnik, Andrés Mauricio Caraballo-Rodríguez, David M. Needham, Corinna Bang, Sören Franzenburg, Marc Bramkamp, Willi Rath, Arne Biastoch, José Carlos Hernández, Ute Hentschel

## Abstract

Development in the sea has long been thought to be a nutritional gamble that disproportionately ends in starvation. Here, we unexpectedly show that components of plastids are incorporated into sea urchin eggs and that these, in turn, benefit developmental fitness. We find chromoplast-derived carotenoid crystals and chromoplast-specific metabolites inside of sea urchin eggs. The light-dependent activity of these chromoplast components influence the subsequent abundance of phytohormones that, in turn, regulate the use of energetic lipids that promote development and survival. Offspring that benefit from these chromoplast components are predicted to disperse further, over larger geographic areas, and use a wider range of currents, including those that cross ocean basins. Data presented here challenge the long-held assumption that components of non-metazoan organelles are unable to enter the germline and be passed between generations. We hypothesize that sea urchins manipulate plastids solely for their self-interest by a process that we term ‘machioplasty,’ with the result of this process being a novel and adaptive form of maternal provisioning.

**SIGNIFICANCE:** Making a fertilizable egg is a complex and carefully regulated process. One long-held assumption is that any component of non-animal organelles are unable to cross the evolutionary valley between non-reproductive and reproductive cells. Here, we unexpectedly show that components of plastids are incorporated into sea urchin eggs and that these, in turn, benefit developmental fitness. Offspring benefit from these plastid components by developing more quickly into larvae and having higher survival due to ability to use phytohormones that influence energetic lipids. This allows offspring to use a wider range of ocean currents, including those that cross entire basins. This challenge the long-held assumption that components of non-metazoan organelles are unable to be passed between generations.

## INTRODUCTION

The most common reproductive strategy amongst marine invertebrates involves producing a high number of small, energy-poor eggs (1, 3). These offspring develop into larvae that acquire nutrients by filtering particles (*e.g.*, phytoplankton, bacteria, and detritus) and taking up dissolved organics from the water column (4–6). Availability of these nutritional resources varies considerably in coastal waters and are limited in the offshore waters where much of development takes place (7, 8). Weeks to months under these conditions amounts to extremely high mortality rates (2, 9, 10), and the few larvae that settle onto the seafloor have often dispersed 10s to 100s of kilometers along the continental slope or across an ocean basin (11, 12). This metabolic puzzle has led others to hypothesize that reproductive output counterbalances these ecological restrictions (1, 3, 13), but a complementary means to bridge this gap may be through microbial symbioses (14, 15).

Animals have a long-standing developmental partnership with microbes (16), and marine invertebrates are no exception. Mothers use their reproductive machinery to transmit microbial symbionts that provide essential amino acids (17), contribute to nutritional plasticity (18), and trigger transitions between life stages (19). Another common feature of these communities is the presence of suspected photosymbionts (20–22), which appear to be particularly common in the eggs, embryos, and larvae of sea urchins (14, 15, 20, 23). Such partnerships could, in principle, enable for long-distance dispersal in waters that are otherwise poorly equip to support this developmental lifestyle (7, 8, 24). Here, we test this hypothesis using *Arbacia lixula*, a sea urchin with an egg that is representative of this major reproductive strategy and whose larvae can disperse across the Atlantic Ocean from well-mixed populations in the Mediterranean Sea and Macaronesia (13, 25).

## RESULTS

### Maternally transmitted plastid structure

We compared the microbial communities of *A. lixula* eggs, embryos, and larvae (that were never fed) using amplicon sequencing of the 16S rRNA gene. These communities were primarily composed of the bacterial phyla Bacteroidota (7.3%), Cyanobacteria (3.4%), Firmicutes (5.8%), and Proteobacteria (80.5%) (Fig. 1A). The majority (78.4% ± 5.0%) of cyanobacterial sequences associated with the egg were, however, unidentified plastid sequences that could be reassigned to photosynthetic eukaryotes (Fig. 1A, 1B). These mother-specific plastid sequences were derived from the Archaeplastida (13.3%), Excavata (14.9%), Hacrobia (45.3%), and Stramenopiles (26.5%) (Fig. 1B, SI Appendix, Fig. S1; SI Appendix, Dataset S1A). We re-analyzed the egg-associated microbiota for a dozen other sea urchin species(23) and found an identical pattern: cyanobacteria and photosynthetic eukaryotes were both present, but photosynthetic eukaryotes represented nearly all of these sequences (SI Appendix, Fig. S2; SI Appendix, Dataset S1B-C).

**Fig. 1:**
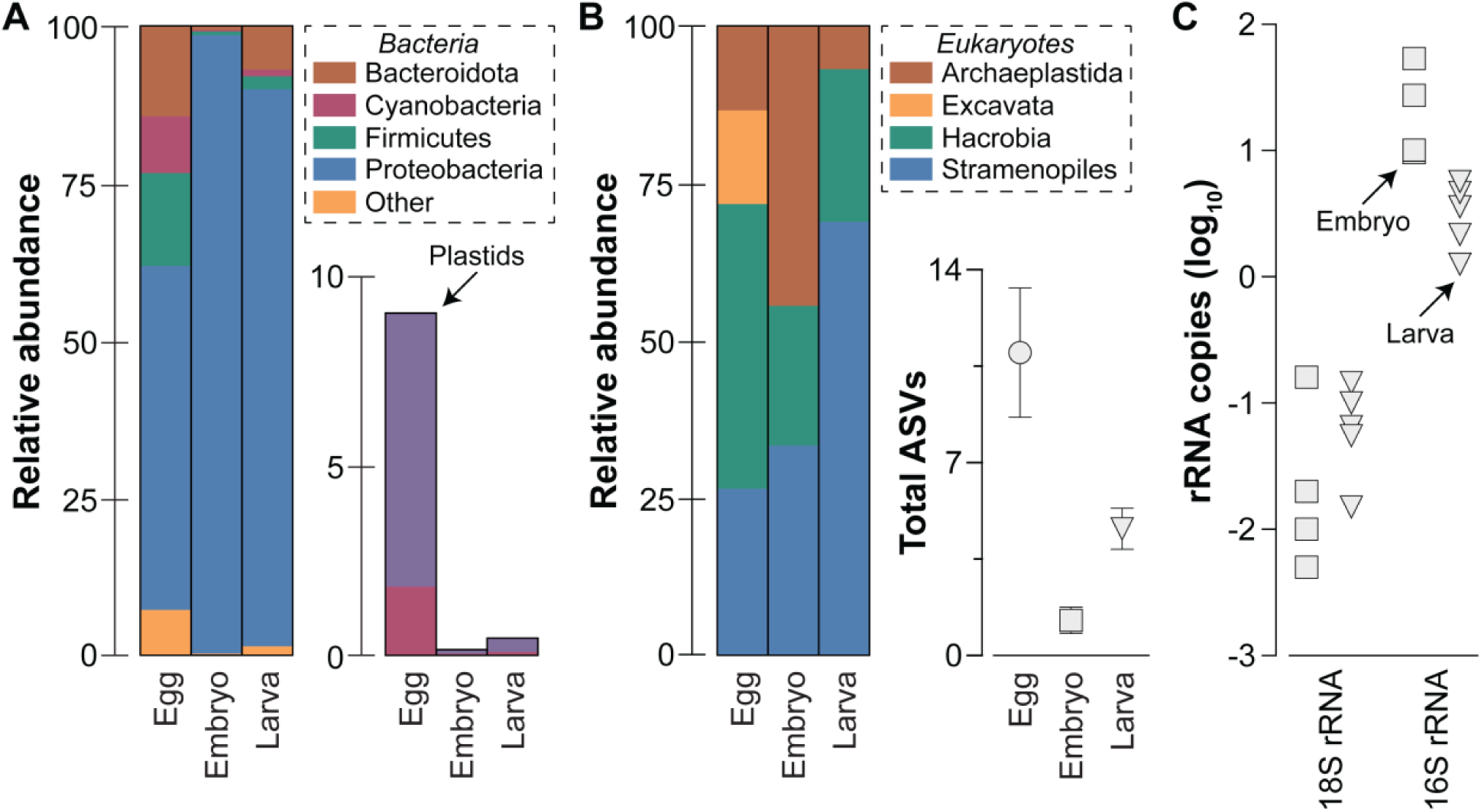
Plastids DNA in sea urchin eggs. (A) One of the main microbial phyla found to associate with the offspring of the sea urchin *Arbacia lixula* were Cyanobacteria, but the majority of these sequences were unidentified plastids. (B) All unidentified plastids could be traced to four major groups and less than a dozen ASVs of photosynthetic eukaryotes. (C) A substantial volume of *A. lixula* eggs would need to be allocated towards photosynthetic eukaryotes if their entire cell was maternally transmitted. A nuclear gene marker (18S rRNA gene) was disproportionately low in abundance in the metagenome of *A. lixula* embryos and larvae compared to a plastid gene marker (16S rRNA gene), implying that mothers may only provide her offspring with plastids.

Plastid sequences were exclusive to and consistently present in the eggs of sea urchins that develop by this major reproductive strategy (SI Appendix, Fig. S3). These plastid taxa (*i.e.*, amplicon sequence variants, ASVs) originated from eight eukaryotic phyla that fall within five kingdoms, with Ochrophyta being the most abundant and the Bacillariophyta (*i.e.*, diatoms) representing ∼90.0% of these sequences (SI Appendix, Fig. S4-5; SI Appendix, Dataset S1D-E). Approximately 91.6% of Ochrophyta ASVs—and ∼74.6% of all ASVs—were derived from diatoms (SI Appendix, Fig. S4-5; SI Appendix, Dataset S1D-E). Sea urchins tend to provide their eggs with a species-specific collection of plastid ASVs [Analysis of Similarity (ANOSIM), p < 0.001], which can more clearly be defined by geography (ANOSIM, p < 0.001) and time (ANOSIM, p<0.001) (SI Appendix, Fig. S6-7; SI Appendix, Dataset S1F-G). Richness of these plastid ASVs directly relates to egg size (linear regression: F_1,109_ = 7.80, p = 0.006, R^2^ = 0.067), whereby sea urchin species with smaller eggs tend to specialize on a few dominant ASVs while those with larger eggs had a relatively even distribution of a dozen or so ASVs (linear regression: F_1,109_ = 68.28, p < 0.0001, R^2^ = 0.385; SI Appendix, Fig. S8-9).

Associating with diverse photosynthetic eukaryotes with cell sizes similar to sea urchin eggs would present a geometric problem for the provisioning of macronutrients that are essential for early development (SI Appendix, Fig. S10) (13, 26). An alternative strategy would be for mothers to provide her eggs with only the plastids from these photosynthetic eukaryotes. We used shotgun metagenomics to quantitatively compare a plastid (16S rRNA) and nuclear (18S rRNA) gene marker for photosynthetic eukaryotes. We found that each *A. lixula* embryo had 25.2 (± 20.7) copies of the 16S rRNA gene and 0.05 (± 0.07) copies of the 18S rRNA gene from photosynthetic eukaryotes (Fig. 1C, SI Appendix, Fig. S11; SI Appendix, Dataset S1H). Notably, zero copies of the 18S rRNA gene were detected for most eukaryotic phyla, while copies of the 16S rRNA gene were consistently present (SI Appendix, Fig. S11; SI Appendix, Dataset S1H). Abundance of the 16S rRNA gene decreased significantly in larvae (7.3×; unpaired t-test, p = 0.006), while the abundance of the 18S rRNA gene remained consistent (unpaired t-test, p = 0.207) (Fig. 1C, SI Appendix, Fig. S4; SI Appendix, Dataset S1C).

We fixed *A. lixula* eggs for fluorescence and transmission electron microscopy to determine whether plastids are maternally transmitted (27). We observed cytoplasmic structures with autofluorescence in most eggs, which lacked chlorophyll but were most consistent for the presence of carotenoids (Fig. 2A, 2B, SI Appendix, Fig. S12). The ultrastructure of these particles were elongated crystals (Fig. 2C, SI Appendix, Fig. S13). The presence of plastid DNA and autofluorescent particles implies that these are most likely chromoplast-derived carotenoid crystals (Fig. 1-2). We could not identify chromoplast-specific starch granules and plastoglobules from similar structures that are commonly found in sea urchin eggs. Carotenoids (*e.g.*, β-carotene and chromoplast-specific xanthophylls) were, however, detected in the metabolome of *A. lixula* eggs (SI Appendix, Dataset S1I-J). The presence of multiple hallmark structures exclusive to chromoplasts (28, 29) implies that mothers provide her eggs with a plastid-derived structure.

**Fig. 2:**
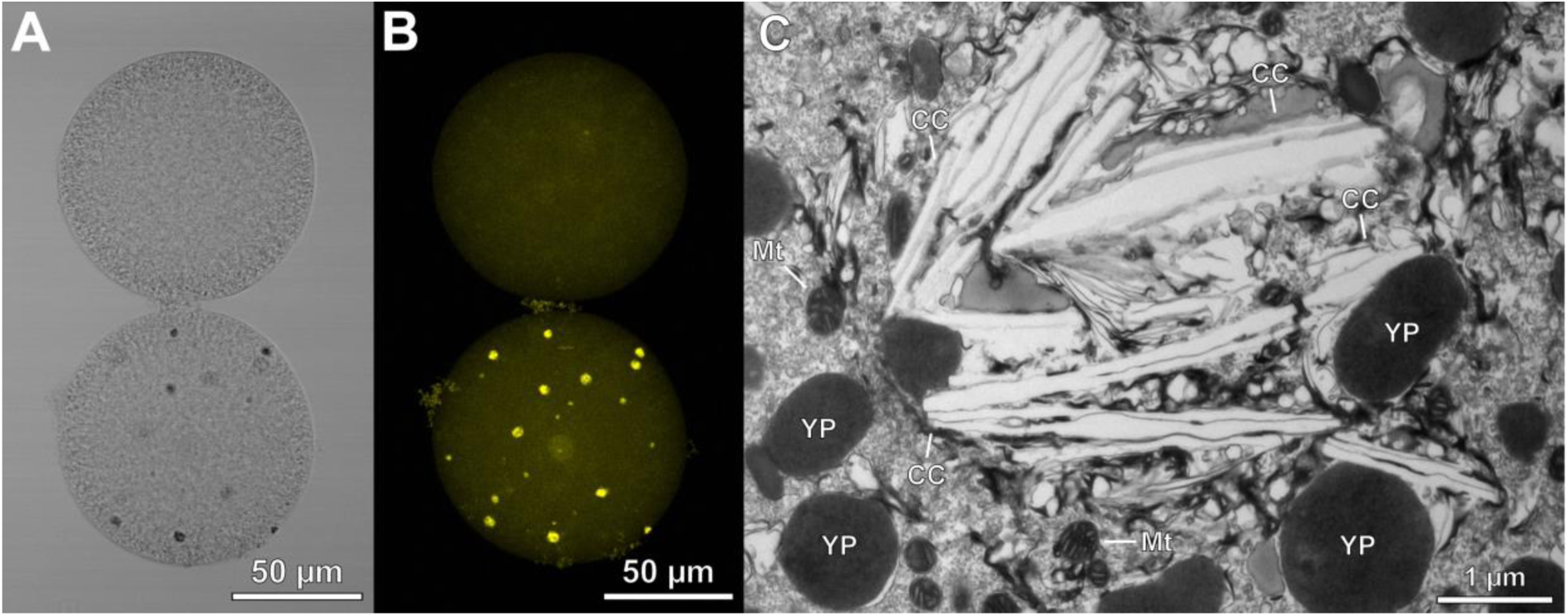
Maternal transmission of a suspected chromoplast structure. (A, B) Micrographs of eggs from the sea urchin *Arbacia lixula* without (top) and with (bottom) cytoplasmic autofluorescent particles. These particles were most consistent for the presence of carotenoids, as shown using (A) transmitted light and (B) fluorescence at an excitation of 561 nm. The light micrograph is a single optical slice, while the florescence micrograph is a maximum projection Z-stack. (C) Cellular structure of these particles are large crystals (CC). Mitochondria (Mt) and yolk platelets (YP) are also noted. The combined fluorescence signature and cellular structure implies that these crystal structures are likely chromoplast-derived carotenoid crystals.

### Benefits to offspring fitness

Chromoplast-derived carotenoid crystals influence plant reproductive fitness and are involved in light-dependent reactions that are independent of photosynthesis (30, 31). Sea urchin development, on the other hand, is not known to be directly impacted by light. If these chromoplast-derived carotenoid crystals are beneficial to *A. lixula* offspring, then a ‘loss of function’ (*i.e.*, darkness) should negatively impact development. We observed that *A. lixula* offspring develop quicker in dark during the initial cell divisions (paired t-test, p = 0.025), but have no observable difference upon gastrulation (paired t-test, p = 1.000; SI Appendix, Fig. S14-15; SI Appendix, Dataset S1K). By four days post-fertilization, 4.8× more offspring developed into 2-arm larvae in light than their siblings in dark (paired t-test, p = 0.026; Fig. 3A, SI Appendix, Fig. S14; SI Appendix, Dataset S1K). The feeding arms of larvae that were cultured in dark grew disproportionately relative to the larval body (unpaired t-test, p = 0.012; Fig. 3B, SI Appendix, Fig. S16-17; SI Appendix, Dataset S1K); thus, larvae in dark expressed the long-arm phenotype that corresponds with a nutritional restriction (32). However, unlike typical morphological plasticity, the gut volume did not differ between *A. lixula* larvae that were cultured in light and dark (unpaired t-test, p = 0.952; Fig. 3B, Extended Data 16-17; SI Appendix, Dataset S1K).

**Fig. 3:**
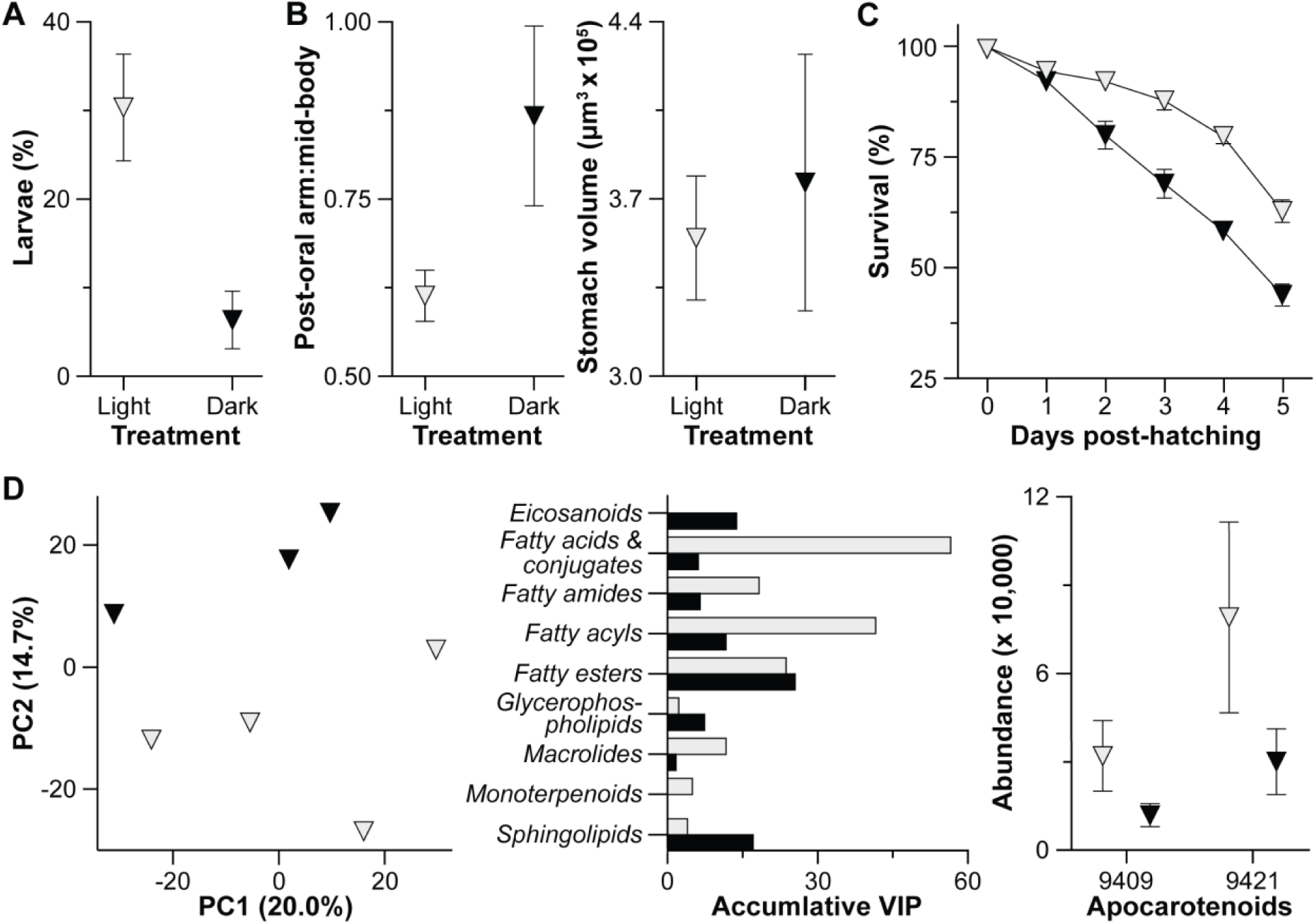
Benefits to offspring fitness. (A) More offspring of the sea urchin *Arbacia lixula* developed into 2-arm larvae in light (gray) than their siblings in dark (black) by 4 days post-fertilization. (B) Feeding arms of larvae that were cultured in dark grew disproportionately to the larval body (left), but did not exhibit a change in gut volume (right); thus, larvae in dark expressed the long-arm phenotype that corresponds with a nutritional limitation. (C) Survival was higher for offspring cultured in light, of which was also affected by time for both treatments as well as the interaction between these factors. Differences in light-induced survival was observed on days two through five. (D) Larvae cultured in light and dark exhibit organism-wide differences in their metabolome (left). This was predominantly driven by a shift in fatty acid metabolism, from the catabolism of energy storage lipids in light to structural lipids in dark (Variable Importance in Projection, VIPs; center). Two of the 717 differentially abundant metabolites are apocarotenoids, which were more abundant for larvae cultured in light (right).

A reduced developmental rate and modification to nutrition are two key factors that directly impact developmental fitness (9), which can be assessed by quantifying survival. We observed that the survival of *A. lixula* offspring was significantly reduced when cultured in dark, as well as with time and the interaction between these factors (two-way ANOVA, time: F_5,54_ = 143.6, p < 0.0001; treatment: F_1,54_ = 109.9, p = 0.0007, interaction: F_5,54_ = 10.31, p < 0.0001; Fig. 3C; SI Appendix, Dataset S1K). Differences in survivorship were observed two days post-hatching (*i.e.*, four days post-fertilization) and in all subsequent days (p < 0.001 for each day), such that survivorship was 1.4× higher in light at five days post-hatching (Fig. 3C; SI Appendix, Dataset S1K).

We then sought to understand how what we presume are chromoplast-derived carotenoid crystals impact the developmental metabolism of *A. lixula*. Light induced an organism-wide divergence in the metabolome during early development (PERMANOVA, eggs: p = 0.181, p < 0.001; Fig. 3D, SI Appendix, Fig. S18; SI Appendix, Dataset S1L). This was predominantly driven by a shift in fatty acid metabolism, from the catabolism of energy storage lipids (*e.g.*, fatty acids, acyls, and esters) in light to structural lipids (*e.g.*, phospholipids and sphingolipids) in dark (Fig. 3D, SI Appendix, Fig. S19; SI Appendix, Dataset S1I-J, S1M). Two of the 717 differentially abundant metabolites were apocarotenoids (*i.e.*, xanthophyll-derived phytohormones that promote growth and development, regulate lipid metabolism, and modulate environmental stress (33–35). These apocarotenoids are a prenol lipid and an uncharacterized apocarotenoid, which were 2.7× more abundant in larvae in light and formed a molecular family with additional phytohormones and an energetic lipid (Fig. 3D, SI Appendix, Fig. S19; SI Appendix, Dataset S1I-J, S1M). We, therefore, find that the incorporation of what we presume are chromoplast-derived carotenoid crystals and their derived metabolites have a broad and integral influence in promoting the developmental fitness of *A. lixula*.

### Enhanced and trans-oceanic dispersal

Development and survival can directly impact dispersal (1–3, 10). We used our survival data to estimate the potential pelagic larval duration for *A. lixula* that do (*i.e.*, light) and do not (*i.e.*, dark) benefit from the light-dependent activity of chromoplast components (Fig. 3C). By quantifying the fecundity of *A. lixula* (SI Appendix, Fig. S3) and deriving the instantaneous mortality rate from our survival data (Fig. 3C) (9), we estimate that offspring that benefit from the light-dependent activity of chromoplast components have a 1.8× longer dispersal potential than those without it (SI Appendix, Fig. S20). We then used the VIKING20X (36) circulation model of the Atlantic Ocean to simulate dispersal. We estimate that offspring that benefit from the light-dependent activity of chromoplast components disperse 1.6× further (paired t-test, p < 0.0001) and across a 4.7× larger geographical area (paired t-test, p < 0.0001) than those without that benefit (Fig. 4A, SI Appendix, Fig. S21). A longer dispersal is estimated to enable 2.9× more larvae additional dispersal paths for a successful settlement (paired t-test, p < 0.001; SI Appendix, Fig. S21-24).

**Fig. 4:**
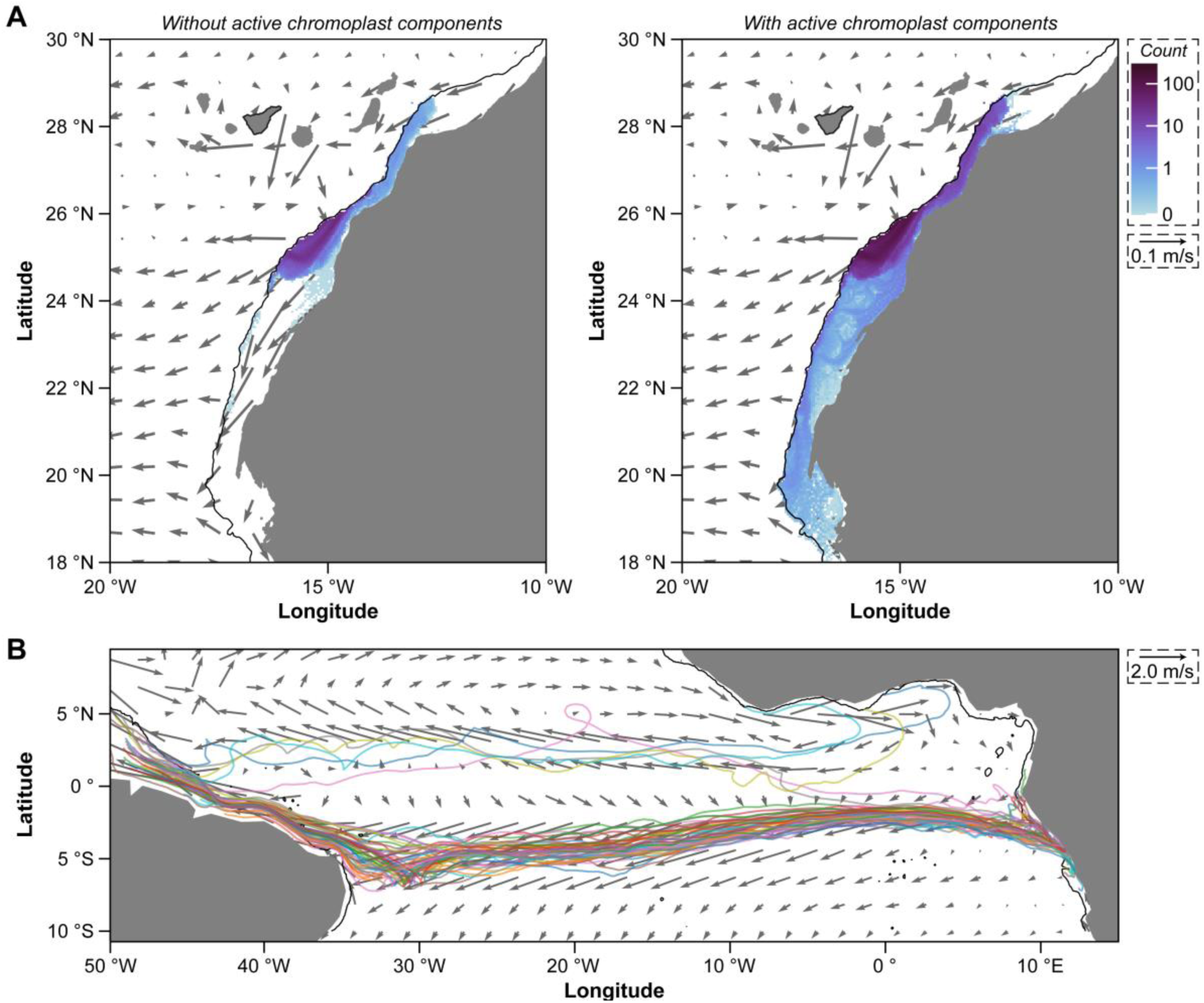
Enhanced and trans-oceanic dispersal. (A) Distribution of particles on the African shelf after 102 (*i.e.*, without the light-dependent activity of chromoplast components; left) and 181 (*i.e.*, with the light-dependent activity of chromoplast components; right) days of being released from Tenerife (black bordered island). Grey arrows show the 10-year mean velocity in the release depth from VIKING20X, with every 10^th^ arrow is shown. Black contours around the African shelf mark the 500 m isobath. (B) Only particles that dispersed for 181 days from the southernmost point of *Arbacia lixula*’s range were predicted to disperse across the Northern Atlantic Ocean from the coast of Africa to Brazil. Fifty example trajectories are shown. Each plot is the accumulation of annual releases over 10 years. Grey arrows as in A, but only every 50^th^ arrow is shown.

Offspring that benefit from the light-dependent activity of chromoplast components also take fewer generations to spread across the longitudinal range of *A. lixula* (SI Appendix, Fig. S25). At the southernmost point of this range, only larvae that benefit from the light-dependent activity of chromoplast components—an estimated 0.3% of the total—are then predicted to disperse across the Atlantic Ocean (SI Appendix, Fig. S26-29). Nearly all of these larvae enter the Angola Current, merge with the South Equatorial Current, and then settle on the northern coast of Brazil (Fig. 4B, SI Appendix, Fig. S26-29). This east to west trans-Atlantic journey is successful each year despite the inter-annual variability in the strength of these currents (SI Appendix, Fig. S28). Once on the northern coast of Brazil, offspring that benefit from the light-dependent activity of chromoplast components could disperse southward along the continental shelf, back to the African coast, and to Cape Verde (SI Appendix, Fig. S30). It, however, would take 2.0× longer to reach the Azores and 4.3× longer to reach the Canary Islands than our estimated pelagic larval duration (SI Appendix, Fig. S31).

## DISCUSSION

Making a fertilizable egg is a complex and carefully regulated process that involves the provisioning of nutrients and maternal RNAs (37, 38). One long-held assumption is that components of non-metazoan organelles (*e.g.*, plastids) are unable to cross the evolutionary valley between non-reproductive (somatic) and reproductive (germline) cells. Sea slugs (39) and flatworm (40) are the only animals known to retain stolen plastids from photosynthetic eukaryotes, yet this highly stable organelle is not transmitted between generations. Data presented here provide initial evidence to challenge this assumption. Specifically, that the sea urchin *A. lixula* maternally transmits multiple components of plastids (*i.e.*, plastid DNA, what we suspect are chromoplast-derived carotenoid crystals, and chromoplast-specific metabolites). Further supporting this hypothesis is that no other crystal structure has been reported in sea urchin eggs despite the cellular biology of fertilization and their pigments having been well-characterized (41, 42) and that sea urchin eggs contain pigments and chloroplast-associated glycolipids that have been collectively referred to as the ‘chloroplast lipid’ peak (43).

If sea urchins maternally transmit chromoplast-derived carotenoid crystals, then it remains paramount to understand the process that encompasses their isolation and incorporation. We hypothesize that this process—preliminarily termed ‘machioplasty’ for being characterized by manipulation, indifference, and self-interest—involves three stages. First, chloroplasts derived from diatoms and other photosynthetic eukaryotes are ingested by adult sea urchins while grazing (39, 40). Second, isolated chloroplasts differentiate into non-photosynthetic chromoplasts during the weeks to months spent inside the dark body cavity (28, 29), a conversion that has been observed in sea slugs (44). Third, entire chromoplasts or, more likely, their specific components are then phagocytosed during oogenesis. We believe that this broad framework allows for the hypothesized process to be characterized and for the direct impact of what we presume are chromoplast-derived carotenoid crystals on developmental fitness to then be quantified.

There is no indication that what we presume are chromoplast-derived carotenoid crystals are within an organelle inside the egg. There is, however, evidence supporting the differential abundance of phytohormones and a light-dependent but photosynthesis-independent reaction that benefits developmental fitness. Light is expected to promote the swimming of these phototaxis larvae that, in turn, should utilize maternal energetics and result in the expression of the phenotype associated with nutrient limitation (45–47). Light, or the lack thereof, is not known to directly influence the development, phenotype, or survival of marine invertebrate larvae that develop by this major reproductive strategy, as we have observed. Chromoplast-derived carotenoid crystals and their derived metabolites are involved in light-dependent reactions in photosynthetic eukaryotes (30, 31). Moreover, experimentally enriching xanthophylls in the diet of adult sea urchins can enhance offspring size, developmental rate, and survival (48). We, thus, presume that these plastid components impact the developmental fitness of *A. lixula*. A benefit to offspring fitness and no clear direct benefit to the adults should, in turn, have a net positive benefit across the entire host’s lifecycle, which may result in this being an adaptive process (49).

Incorporating what we presume are chromoplast-derived carotenoid crystals should increase the total resources provisioned into an egg and, in turn, produce a more fit offspring. This additional maternal resource enhances dispersal potential, which is most pronounced through the prediction that only offspring that benefit from the light-dependent activity of these chromoplast components may undergo trans-Atlantic dispersal. The metabolic benefit of what we presume are chromoplast-derived carotenoid crystals could provide offspring with the potential to cross an ocean basin and use this major oceanic current as a highway for gene flow (12, 25, 50). Providing offspring with plastid-derived components in the sunlit ocean could, thus, serve as a mechanism to promote long-distance (*i.e.*, teleplanic) dispersal. This developmental feature is observed for most groups of marine invertebrates and in every major ocean, including sea urchin larvae that use the South Equatorial Current to cross the Atlantic Ocean (12, 50). These plastid-derived components are hypothesized to be a novel form of maternal provisioning and a complementary metabolic resource to the nutrients acquired by filtering particles, taking up of dissolved organics, or through microbial symbioses (4–6, 17).

## MATERIALS AND METHODS

*Specimen collection and offspring culturing*. Adult *A. lixula* were collected from Radazul Beach (Tenerife, Canary Islands, Spain) during 2021 to 2024, transferred dry to the University of La Laguna, and maintained in ambient seawater. Adults were spawned, fertilized, and full-sibling replicates were cultured according to Strathmann (51). Each culture was diluted to 2 embryos/mL and each male-female pair was divided amongst separate light and dark beakers.

*Sample collection, extraction, and sequencing*. We collected eggs, embryos, and larvae from each ‘light’ beaker for amplicon and metagenomic sequencing. Offspring at each stage were collected from each male-female, and were concentrated using a microcentrifuge, removed of seawater, and preserved for long-term storage. Total DNA was extracted from all samples and DNA kit blanks. Total DNA was then quantified and diluted to 0.5 ng/µL for amplicon and not diluted for metagenomics. Libraries were prepared for amplicon (V3/V4 region of the 16S rRNA gene) and metagenomics and sequenced by Illumina MiSeq (v3, 2×300 bp paired-end reads) and Illumina NovaSeq (S4, 2×150 bp paired-end reads), respectively.

*Amplicon of A. lixula*. Raw reads were imported into QIIME 2 (52), where they were paired, filtered by quality score, and denoised. Features were analyzed as amplicon sequence variants (ASVs) and assigned taxonomy using SILVA (53). Archaea, contaminants, and low-read samples were discarded. The filtered data table was rarified. We summarized taxonomic groups of bacteria for each sample, and then filtered this data table to only include “o Chloroplast” for taxonomic reassigned using PhytoREF (54).

*Amplicon of sea urchins*. We compiled all published 16S rRNA datasets for sea urchin eggs. We generated taxonomic profiles for each sea urchin and then compared the relative abundance of “o Chloroplast” to the Cyanobacteria, and then reassigned taxonomy for “o Chloroplast” using PhytoREF (54). We transformed the plastid-only data table to relative frequency to avoid discarding invaluable sequences. We then generated a phylogenetic tree for all plastid ASVs, and manually curated the prevalence of plastids in each sample and ASVs across species. We calculated Jaccard distances and visualized these using principal coordinate analyses to compare the community composition across host species, geography, and years. We then compared host phylogeny to a plastid dendrogram grouped by host species. Lastly, we assessed four measures of alpha diversity and compared these to sea urchin life-history traits.

*Metagenomics*. Raw reads were processed for quality and host sequences within MetaWRAP (55). rRNA gene sequences were identified using METAXA2 (56) and their taxonomy was assigned using the same process described above. Plastid (16S rRNA gene) and nuclear sequences (18S rRNA gene) were quantified, and all equivalent groups of photosynthetic eukaryotes were then compared.

*Microscopy*. Eggs were fixed for florescence and transmission electron microscopy. In the former, eggs were assessed for autofluorescence above 620 nm at excitation wavelengths of 405 nm, 488 nm, 561 nm, and 640 nm. Maximum intensity projections of Z-stacks were generated, while single Z-planes were used for transmitted light images. In the latter, samples were embedded in resin, dehydrated, and infiltrated with Epon and sectioned using an ultramicrotome. Grids were inspected for particles that could be autofluorescence and appear non-native to sea urchins.

*Development, morphology, and survival*. We measured egg size, development, and morphology as part of the experiment outlined in ‘*Sample collection, extraction, and sequencing*.’ Egg diameter was measured for ‘light’ beakers, while developmental stages at 0 hours post-fertilization (hpf), 4 hpf, 18 hpf, 2 dpf, and 4 dpf were assessed for all beakers. Offspring were staged by developmental stage and morphometrics were performed for 2-arm larvae at 4 dpf.

Survivorship was quantified in separate experiments using the same experimental design. Offspring of male-female pairs developed until hatching, after which each culture was diluted to 2 embryos/mL. Embryo density was counted daily for five days (*i.e.,* until 7 dpf). Five biological replicates were performed in 0.22 μm FSW and the other five were performed in 5.0 μm FSW as a cross-validation for maternal transmission. Both survival datasets were merged because the transmission mechanism was maternal and because the survival rates were statistically identical.

*Metabolomics*. Metabolomics was used in a separate experiment using the exact same experimental design. Samples were prepared for untargeted liquid chromatography tandem mass spectrometry. Raw data files were processed in MZmine (57). Annotations and chemical classification of detected molecules were obtained with SIRIUS (58), which were used for a feature-based molecular network in GNPS2 (59). Metabolite composition was compared using a principal component analysis and a Partial Least Squares Discriminant Analysis to identify differentially abundant metabolites. We summarized the total contribution of these differentially abundant metabolites at the pathway, superclass, and class levels. Lastly, we used normalized data to summarize the abundance of two differentially abundant metabolites and assessed which metabolites are within their respective molecular family from the feature-based molecular network.

*Estimating and modeling dispersal*. We calculated the duration required for the survival of the last larva from the clutch of *A. lixula* with and without the light-dependent activity of what we presume are chromoplast components. Offspring abundance (N_t_) after a specific time interval (t) was calculated using the instantaneous rate model: N_t_ = N_0_ e ^-Mt^, where N_0_ is fecundity, e is the Naperian constant, and M is the instantaneous rate of mortality (9). M was calculated using the survival data for *A. lixula* offspring. Fecundity (N_0_) was estimated from individuals at Radazul Beach.

We integrated these specific time intervals into VIKING20X (36), an oceanographic model of the Atlantic Ocean. We used the Lagrangian simulation software tool Parcels (60) to simulate the 3D advection of virtual larvae in the upper vertical levels from 2007 to 2017. Virtual particles (10,000 per week for 10 years) with a lifetime of 102 or 181 days were released in the coastal waters around Tenerife. The abundance, geographical location, and distance of particles from each treatment that reached the continental shelf of Africa were tracked. Using this framework, we determined how many generations it would take for virtual larvae to spread along the African shelf and reach the southernmost range for *A. lixula*. We then determined if and, if so, which locations could be used as the departure area for trans-Atlantic dispersal, and if those particles could then disperse to back to Macaronesia and towards Uruguay. An additional modeling experiment was performed to determine when virtual particles reached the Azores and the Canary Islands.

### Data and code availability

Raw amplicon and metagenomic sequencing files have been deposited to the Sequence Read Archive of the NCBI under the BioProject accession number PRJNA1170906. Raw metabolomic data and the complete methodological details for the LC-MS/MS have been deposited to the MassIVE repository (https://massive.ucsd.edu/) under the accession number MSV000097754. The FBMN analysis can be accessed directly using the following link: https://gnps2.org/status?task=31e1342caaa14a649633c328148474c1. Code corresponding to the amplicon, metagenomic, and metabolomic datasets are all available on GitHub (https://github.com/TylerJCarrier/Arbacia_DevelopmentSymbiosis). Code used to simulate dispersal are available on a separate GitHub (https://github.com/geomar-od-lagrange/2023_sea_urchin_tenerife).

## Acknowledgements

We thank members of the Marine Community Ecology and Conservation lab at the University of La Laguna (Tenerife, Spain) as well as the Research Unit Marine Symbioses at the GEOMAR Helmholtz Centre for Ocean Research (Kiel, Germany) for assistance with field collections and logistics. We also thank many colleagues for fundamental conversations and the croquetas throughout this endeavor. Samples were collected and exported in coordination with the Spanish Government (SGBTM/BDM/AUTSPP/70/2022 and 1488098). Authorization under the Nagoya Protocol was not required for this work because it fell under the definition of “exclusively taxonomic purposes” and, thus, it did not constitute a utilization of genetic resources under the Spanish access regulation. This study was funded by the Alexander von Humboldt Foundation, German Research Foundation (project B1 of the CRC 1182 “Origin and Function of Metaorganisms”; Project ID: 261376515), GEOMAR Helmholtz Centre for Ocean Research, Spanish Ministry for Science and Innovation (PGC2018-100735-B-I00; MCIU/AEI/FEDER, UE), and the Symbiosis in Aquatic Systems Initiative of the Gordon and Betty Moore Foundation (GBMF12120). Microbiome sequencing received infrastructure support from the German Research Foundation German Research Foundation [DFG Excellence Cluster 2167 “Precision Medicine in Chronic Inflammation,” DFG Research Unit 5042 “miTarget,” and the Next Generation Sequencing Competence Network (Project IDs: 423957469 and 407495230)].

## Author contributions

T.J.C. designed this study. T.J.C., A.R.-N., and J.C.H. performed field collections and conducted organismal experiments. T.J.C., A.R.-N., D.M.N., U.H. and C.B. generated and analyzed amplicon sequencing data. T.J.C. performed the amplicon meta-analysis. T.J.C., D.M.N., and S.F. generated and analyzed metagenomic data. U.R., T.J.C., U.H. and M.B. performed and interpreted the microscopy. M.C. and T.J.C. generated and analyzed the metabolomics data. T.K., W.R., A.B., and T.J.C. designed and performed the biophysical model. T.J.C. wrote the manuscript with contributions from all co-authors.

