## Appendix for "Maternal transmission of a plastid structure enhances offspring fitness"

This PDF file includes:

- Supplementary methods
- Fig. S1 to S33
- Dataset S1A to S1N
- Supplemental references

### MATERIALS AND METHODS

#### *Specimen collection*

Adult *Arbacia lixula* were collected by snorkel from Radazul Beach (Tenerife, Canary Islands, Spain; 28.401487, -16.324368) from 2021 to 2024. Adult sea urchins were transferred dry to the University of La Laguna within one hour, where they were maintained in an aerated aquarium with seawater from Radazul Beach until spawning a couple hours later.

#### *Offspring culturing*

Gravid adults were spawned by an intracoelomic injection of 0.50 M KCl. Females were spawned into 5.0 µm filtered seawater (FSW) from Radazul Beach and males were spawned dry and stored on ice. Female and male gametes were collected to generate full-sibling replicates (n = 5) by following Strathmann (1). Briefly, a few drops of diluted (1:1000) sperm from the male pair was added to the total clutch of the female pair. Egg and sperm were gently mixed. Fertilization was assessed 30 minutes later using a compound microscope (Leica DM1000 with a Canon EOS 1200D camera). Cultures were diluted to 2 embryos/mL and each male-female pair was divided amongst separate light and dark beakers. Dark beakers were covered in tinfoil, equipped with a cardboard lid with an overhanging layer of tinfoil, and a loose-fit black garbage bag on the stirring paddle. All beakers were stirred gently and >90% of the FSW was replenished every other day. Eggs, embryos, and larvae were cultured in ambient FSW at room temperature and were never provided phytoplankton or another external dietary source.

#### *Sample collection, extraction, and sequencing*

We collected eggs (0 hours post-fertilization, hpf), embryos (blastula; 18 hpf), and early-stage larvae (4 days post-fertilization, dpf) from each 'light' beaker (n = 5) for amplicon and metagenomic sequencing; both sequencing types were from the same sample. Approximately 200 offspring were collected from each male-female pair for each of the developmental stages. Offspring were concentrated using a microcentrifuge, the seawater was removed with a sterile pipette, and tissues were preserved in 250 µL of 70% ethanol for long-term storage at -20 °C. Samples were then transferred on blue ice to the GEOMAR Helmholtz Centre for Ocean Research (Kiel, Germany).

Total DNA was extracted from all samples and DNA kit blanks (n = 3) according to the manufacture's protocol for the DNeasy® Blood & Tissue Mini Kit (Qiagen). Total DNA was then quantified using the dsDNA HS Assay Kits for the Qubit Fluorometer (Thermo Fisher Scientific) following the manufacture's protocol. Samples were diluted to 0.5 ng/µL for amplicon sequencing and were not diluted for metagenomic sequencing. Samples for amplicon (V3/V4 region of the 16S rRNA gene) and metagenomics were sent to the Competence Center for Genomic Analysis (Christian-Albrechts University of Kiel, Germany) for library preparation and sequencing by Illumina MiSeq (v3, 2x300 bp paired-end reads) and Illumina NovaSeq (S4, 2x150 bp paired-end reads), respectively.

#### *Amplicon analysis of A. lixula*

Raw amplicon reads with quality information were imported into QIIME 2 (v. 2022.11; 2), where forward and reverse reads were paired using VSEARCH (3), filtered by quality score, and denoised using Deblur (4). QIIME 2-generated 'features' were analyzed as amplicon sequence variants (ASVs; 5) and were assigned taxonomy using SILVA (v. 138; 6). Sequences matching to Archaea, those that were disproportionately present in the DNA kit blanks, and samples with less than 3,000 reads were discarded. The filtered data table was then rarified to 3,482 sequences (SI Appendix, Fig. S32). We summarized taxonomic groups of bacteria for each sample from the SILVA (6) assignment. We then filtered this data table and used PhytoREF (7)—a database for plastidial 16S rRNA gene sequences of photosynthetic eukaryotes—to re-assign taxonomy for all ASVs that were previously assigned to "o__Chloroplast." Taxonomic overlap in the plastid ASVs that were present in the eggs of each female *A. lixula* was determined using InteractiVenn (8).

*Amplicon meta-analysis of sea urchins*

We compiled all published 16S rRNA datasets for sea urchin eggs (SI Appendix, Dataset S1B) (9-12), as well as single samples of *Eucidaris thouarsii* and *Eu. tribuloides* that was collected during, but not published with, Carrier *et al.* (10). We generated taxonomic profiles for each sea urchin species and then tallied the relative abundance of "o__Chloroplast" to the non-plastid Cyanobacteria sequences, and then reassigned the taxonomy for "o__Chloroplast" using PhytoREF (7). We transformed the plastid-only data table to relative frequency to avoid discarding invaluable sequences. Using this, we generated a phylogenetic tree for all plastid ASVs within QIIME 2 (2) using MAFFT(13) and FastTree(14). This tree was then visualized using the Interactive Tree Of Life (15) and stylized using Adobe Illustrator (v. 24.0.1). Mean relative frequency of each plastid ASV was added as metadata to the phylogenetic tree. Next, we manually curated the prevalence of plastids in each sample and ASVs across species. We then calculated Jaccard distances for all samples, visualized these using principal coordinate analyses in Prism (v. 9.0.0), and stylized these using Adobe Illustrator. ANOSIM and PERMDISP, as well as their respective pairwise comparisons, were performed within QIIME 2 (2) to test whether community composition and dispersion varied between host species, geography, and years.

Jaccard distances were then used to construct a dendrogram within QIIME 2 (2). The data table for this analysis was collapsed by host species and then was rarified to 447 sequences; this was the only rarified analysis. We then tested for phylosymbiosis by comparing topological congruence with a Cytochrome C Oxidase Subunit I (COI) gene tree using sequences from the National Center for Biotechnology Information (NCBI) (SI Appendix, Dataset S1B). These sequences were imported into the Molecular Evolutionary Genetics Analysis software (v. 11.0.9; 16), aligned with MUSCLE (17), and trimmed. This relationship was inferred using maximum likelihood with the optimized DNA substitution model (GTR+G+I, as determined by BIC criteria) and 1,000 bootstrap replicates. Phylosymbiosis was tested using the Robinson-Foulds metric in TreeCmp (v. 2.0; 18) and matching cluster metrics with 10,000 random trees (19). Lastly, four measures of alpha diversity (total ASVs, Faith's phylogenetic distance, McIntosh evenness, and McIntosh dominance) were calculated for all samples. These were compared across species using one-way ANOVAs and Tukey's post-hoc tests for pairwise comparisons. Total ASV estimates were then compared to published values for egg diameter (20-23), total energetic content (21, 24-26), and pelagic larval duration (20, 27-32) for these sea urchin species using separate linear regression analyses.

*Metagenomic analysis*

Raw metagenomic reads were quality filtered, trimmed, and removed of host sequences within MetaWRAP (33). The *S. purpuratus* genome (v. 5.0) from Echinobase (34, 35) was used to minimize host reads because there was no published genome for *A. lixula* at the time of this analysis. Furthermore, we removed mitochondrial sequences using published mitochondrial genomes for *A. lixula* (NC_001770), *Echinometra mathaei* (NC_034767), *Hemicentrotus* *pulcherrimus* (NC_023771), *Heterocentrotus mammillatus* (NC_034768), and *Lytechinus variegatus* (NC_037785) from NCBI.

rRNA gene sequences were identified in the forward sequences using METAXA2 (36). All rRNA gene sequences were imported into QIIME 2 (2) for taxonomic assignment using the default parameters of the VSEARCH (3) consensus classifier. Plastid sequences (*i.e.*, 16S rRNA gene) were identified using PhytoREF, while nuclear sequences (*i.e.*, 18S rRNA gene) were identified using the combined Protist Ribosomal Reference ($PR^2$) ((37)/EukRef (38) 18S rRNA databases. The quantity of 16S and 18S rRNA gene sequences were counted for all equivalent groups of photosynthetic eukaryotes (SI Appendix, Dataset S1H). Sequences for 16S and 18S rRNA gene were then summed separately and compared between developmental stages using unpaired two-tailed t-tests within Prism.

*Microscopy*

Florescence microscopy was used to assess whether there were autofluorescent signatures consistent with plastid pigments in *A. lixula* eggs. Eggs (n = 10) were fixed at room temperature in 4% formaldehyde buffered in ambient seawater supplemented with 20 mM HEPES. This fixative was replaced after 1 hour with 1% formaldehyde buffered in ambient seawater with 20 mM HEPES buffer, before being stored long-term at 4 ºC. Eggs were assessed for autofluorescence above 620 nm at excitation wavelengths of 405 nm, 488 nm, 561 nm, and 640 nm using an LSM 900 with Airyscan microscope (Zeiss) in the Multiplex mode. Maximum intensity projections of Z-stacks were generated using the ZEN Blue (v. 3.2) software (Zeiss). Transmitted light images at single Z-planes were acquired with the LSM mode. A Plan-Apochromat 20x (0.8 NA) objective was used for all applications.

Transmission electron microscopy was used to assess the ultrastructure of *A. lixula* eggs from the same females (n = 10) that were sampled for fluorescence analysis. Eggs were fixed with 0.5% glutaraldehyde buffered in ambient seawater for 1 hour and then stored long-term at 4 ºC (39). For resin embedding, the eggs were washed with deionized water and glycine to remove the fixative, post-fixed with 1% osmium tetroxide in 1.5% potassium ferricyanide for 1 hour at room temperature, washed several times with deionized water, treated with 0.1% tannic acid in 50 mM HEPES (pH 7) for 3 minutes at room temperature, with 2% aqueous uranyl acetate for 1 hour, and then washed again with deionized water. Samples were dehydrated using a graded ethanol series, followed by 100% acetone, and then progressively infiltrated with Epon™ 812 substitute resin (Sigma-Aldrich) and heat polymerized. Ultrathin 80-nm sections, prepared using the UC7 ultramicrotome (Leica) and diamond knives (Diatome), were contrasted with saturated aqueous uranyl acetate and lead citrate. Grids were inspected in a Tecnai G2 Spirit BioTWIN transmission electron microscope using a side-entry CCD MegaView III G2 camera and iTEM software (v. 5.0) (Olympus Soft Imaging Solutions).

*Development, morphology, and survival*

We measured egg size, development, and morphology as part of the experiment outlined in '*Sample collection, extraction, and sequencing*.' Egg diameter (n = 10 per replicate) was measured at 0 hpf for all 'light' beakers, while the developmental stage (n = 25 per replicate per timepoint) at 0 hpf, 4 hpf, 18 hpf, 2 dpf, and 4 dpf was assessed for all beakers. Offspring were staged as either egg, 2-cell embryo, 4-cell embryo, 8-cell embryo, 16-cell embryo, blastula, gastrula, prism-stage larva, or 2-arm larva. Morphometrics (*i.e.*, mid-body line, post-oral arms, stomach height, and stomach width) were performed on all 2-arm larva at 4 dpf using ImageJ (v.

1.9.2; 40). A paired two-tailed t-test was performed within Prism to test whether development was similar in light and dark conditions. Due to a limited number of 2-arm larvae in dark, unpaired two-tailed t-tests were performed within Prism for each morphological measurement to test whether larval morphology was similar in light and dark conditions.

Survivorship was quantified in separate experiments using the exact same experimental design. Offspring from male-female pairs (n = 10) developed until hatching (*i.e.,* 2 dpf), after which triplicate counts of each culture density was performed and were subsequently diluted to 2 embryos/mL in 1 L of FSW. Cultures were reversed filtered to 100 mL each day, the number of embryos were counted in triplicate 1 mL aliquots using a dissecting microscope (Leica EZ4), and then returned to the beaker along with fresh FSW. This was performed for five days (*i.e.,* until 7 dpf). Five of the biological replicates were performed in 0.22 µm FSW and the other five were performed in 5.0 µm FSW as a cross-validation for maternal transmission. Both survival datasets were merged because the transmission mechanism was maternal and because the survival rates were statistically identical between both conditions (two-way ANOVA for light, time: $F_{5,40}$ = 102.8, $p < 0.0001$; treatment: $F_{1\,8}$ = 0.04, =, interaction: $F_{5,40}$ = 3.91, $p < 0.0055$; two-way ANOVA for dark, time: $F_{5,40}$ = 93.33, $p < 0.0001$; treatment: $F_{1\,8}$ = 0.38, p = 0.554, interaction: $F_{5,40}$ = 1.06, p = 0.399; SI Appendix, Dataset S1K). A two-way ANOVA was performed within Prism to test whether survivorship differed between treatments and over time/development.

*Metabolomics*

Metabolomics was used in a separate experiment using the exact same experimental design as development and survival. Specifically, the clutch of spawned females (n = 5) was equally distributed between separate 1 L beakers for light and dark. Three technical replicates of ~1,000 eggs were collected from all beakers, which were immediately centrifuged to concentrate the tissues, the 0.22 µm FSW was removed, samples were flash frozen using liquid nitrogen, and then transferred to a -80 °C freezer for long-term storage. The remaining eggs were fertilized with dilute sperm to create full-sibling replicates. Excess sperm was removed 1 hour later by multiple 95% water changes and fertilized embryo were then provided two days to develop. Hatched blastulae were isolated, the cultures were diluted to 10 embryos/mL following multiple 95% water changes, and the cultures were then provided 3 additional days to develop (*i.e.*, when there was a maximal differential in survival; Fig 3C). At 5 dpf, three technical replicates of ~1,000 offspring were collected from all beakers and processed as described above. Four and three biological replicates could be collected from the light and dark treatments of larvae, respectively. Blank tubes and 0.22 µm FSW seawater were also collected as controls (n = 5 each). All samples were then shipped on dry ice to the University of California, San Diego (CA, USA).

Samples were prepared for liquid chromatography tandem mass spectrometry (LC-MS/MS) according to an established protocol (41). Untargeted LC-MS/MS was then performed using a Vanquish Flex Ultra-High-Performance Liquid Chromatography (UHPLC) system that was coupled with an Orbitrap Exploris™ 240 Mass Spectrometer (Thermo Fisher Scientific). Chromatographic separation was performed using a Kinetex® 2.6 µm Polar C18 reversed phase UHPLC column with a constant flow rate of 500 µL/min (Phenomenex). LC-MS/MS acquisition followed an isocratic elution and a multistep linear gradient (SI Appendix, Dataset S1N). Data dependent acquisition mode was then used for acquisition of tandem MS (SI Appendix, Dataset S1N). Analytical blanks, pool samples, and mixtures of sulfamethazine, sulfamethizole, sulfachloropyridazine, sulfadimethoxine, amitriptyline, and coumarin-314 at 10 µM were injected after every 10 samples as quality control for monitoring instrument performance.

Raw LC-MS/MS data files were processed in MZmine (v. 4.6.1; 42) to generate a feature table, peak list, and batch file of all settings, with peak area of detected ions being used as metabolite abundance (SI Appendix, Dataset S1J). Annotations and chemical classification of detected molecules were obtained with SIRIUS (v. 6.1.1.1; 43) (SI Appendix, Dataset S1I), with these annotations corresponding to level 3 of the Metabolomics Standards Initiative (44, 45). This was then used for a feature-based molecular network (46) in GNPS2 (47). We generated a rarefaction curve to assess whether the metabolite diversity is fully represented (SI Appendix, Fig. S33). We performed a principal component analysis and separate PERMANOVA in R to compare the metabolome across developmental stages as well as between light and dark for eggs and larvae. We then performed a Partial Least Squares Discriminant Analysis to identify differentially abundant metabolites (*i.e.*, Variable Importance in Projection, or VIPs) of larvae in light and dark. We summarized the total contribution of these differentially abundant metabolites in light and dark at the pathway, superclass, and class levels. Lastly, we used normalized data to summarize the abundance of two VIPs and assessed which metabolites are within their respective molecular family from the feature-based molecular network (48).

*Estimating and modeling dispersal*

We calculated the duration required for the survival of the last larva from the clutch of *A. lixula* with and without the light-dependent activity of what we presume are chromoplast components. The abundance of *A. lixula* offspring ($N_t$) after a specific time interval (t; days) can be calculated using the instantaneous rate model: $N_t = N_0 e^{-Mt}$, where $N_0$ is the fecundity, e is the Naperian constant, and M is the instantaneous rate of mortality ($day^{-1}$) (49). M was calculated using the survival data for *A. lixula* offspring in light and dark. Fecundity ($N_0$) was estimated by spawning individuals (n = 5) until eggs were no longer released, standardizing by total volume, performing two subsequent 1:100 dilutions for 1 mL fully concentrated eggs, and counting triplicate 0.5 mL aliquots of dilute eggs using a dissecting microscope (Leica EZ4). The specific time interval was then calculated for offspring that do and do not benefit from the light-dependent activity of chromoplast components.

We integrated these specific time intervals into the VIKING20X (50), a full-scale, nested, and eddy-rich oceanographic model of the Atlantic Ocean. Specifically, we used the VIKING20X-OMIP (50) to perform our dispersal experiments because of its realistic ability to simulate atmospheric forcing (51), to accurately simulate the large-scale horizontal circulation, distribution of mesoscale, overflow, and convective processes, and the representation of regional current systems in the North and South Atlantic Ocean (52, 53). This combination of oceanographic features has previously enabled the VIKING20X to be used to simulate larval dispersal of costal and deep-sea marine invertebrates (54, 55). We used the Lagrangian simulation software tool Parcels (v. 2.3.1; 56) to simulate the 3D advection of virtual larvae based on the daily-mean velocities for ocean currents in the upper 12 vertical levels (~0-130 m) from 2007 to 2017. Virtual particles were released at depths between 0-30 meters in the coastal waters around Tenerife (SI Appendix, Fig. S21). Our simulation released 10,000 particles each week for 10 years (*i.e.*, from 2007 to 2017) to account for seasonal and inter-annual variability. Virtual particles had a lifetime of either 102 (*i.e.*, without the light-dependent activity of chromoplast components) or 181 days (*i.e.*, with light-dependent activity of chromoplast components). The abundance, geographical location, and distance of particles from each treatment that reached the continental shelf of Africa were tracked. Particles that reached the continental shelf of Africa in 25 days or less were not counted due to the minimal pelagic larval duration for *A. lixula* (23). Each of these quantitative assessments were compared statistically using paired two-tailed t-tests within Prism.

Using this same oceanographic framework, we determined how many iterations (*i.e.*, generations) it took for virtual larvae to spread along the African shelf and reach the southernmost range for *A. lixula*. Generation 0 was considered Tenerife and Generation 1 was the resulting dispersal. The location and abundance of particles from Generation 1 were then used to determine the origins and then spread in Generation 2. This was repeated until a particle reached the southernmost range for *A. lixula*. Next, we evenly distributed 2.0 million particles along the African shelf (from 28 °N to 10 °S; SI Appendix, Fig. S25) to determine if any and which locations could be used as the departure area for trans-Atlantic dispersal. Only particles with a pelagic dispersal potential reflecting the benefits from the light-dependent activity of chromoplast components and initially located at the southernmost area could disperse across the Atlantic Ocean (Fig. 4B), so we then tracked paths that these particles took and compared them to known oceanographic currents. Lastly, we evenly distributed 1.6 million particles along the northern coast of Brazil (from 5°S to 10°S; SI Appendix, Fig. S30) to determine if larvae that benefit from the light-dependent activity of chromoplast components could disperse to back to Macaronesia and towards Uruguay (*i.e.*, the southernmost range for *A. lixula* in South America) (57, 58). This modeling experiment was performed for four years to determine when virtual particles reached the Azores and the Canary Islands.

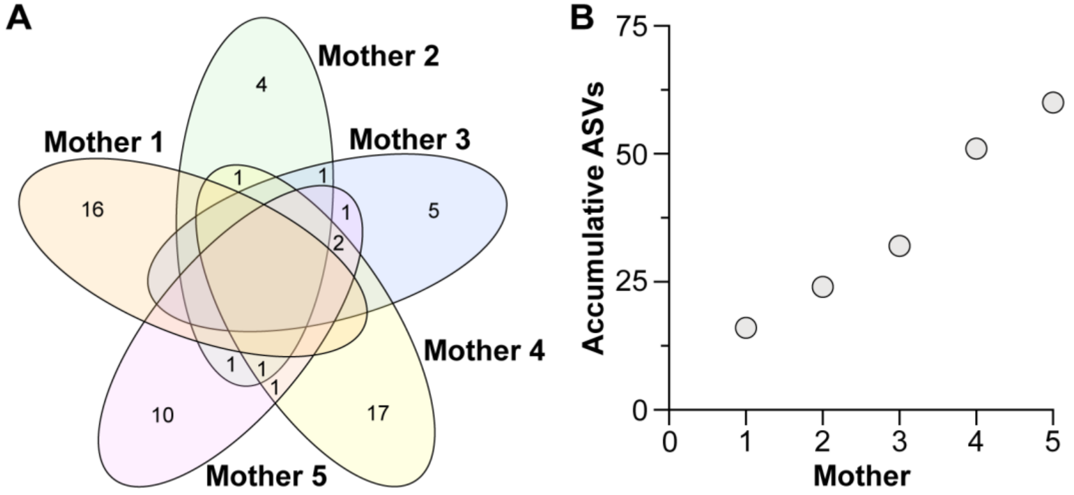


**Fig. S1:** Plastid ASVs in sea urchin eggs. Minimal taxonomic overlap (A) and the diversity (B) of plastid ASVs in egg from different individuals of the sea urchin *Arbacia lixula*. Empty cells in the Venn diagram have zero ASVs.

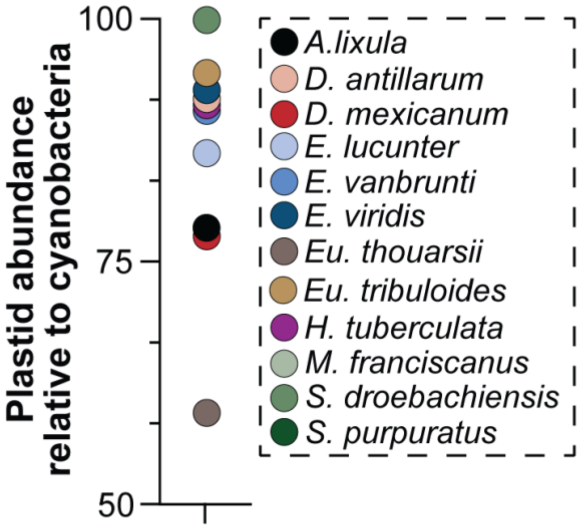


**Fig. S2:** Plastid abundance relative to cyanobacteria. Abundance of plastid sequences from photosynthetic eukaryotes relative to that of cyanobacteria in the egg-associated microbiota of the twelve sea urchins that have been studied to date. Note: *Heliocidaris erythrogramma* that is present in Fig. S3 was not included here because these taxonomic groups are not present.

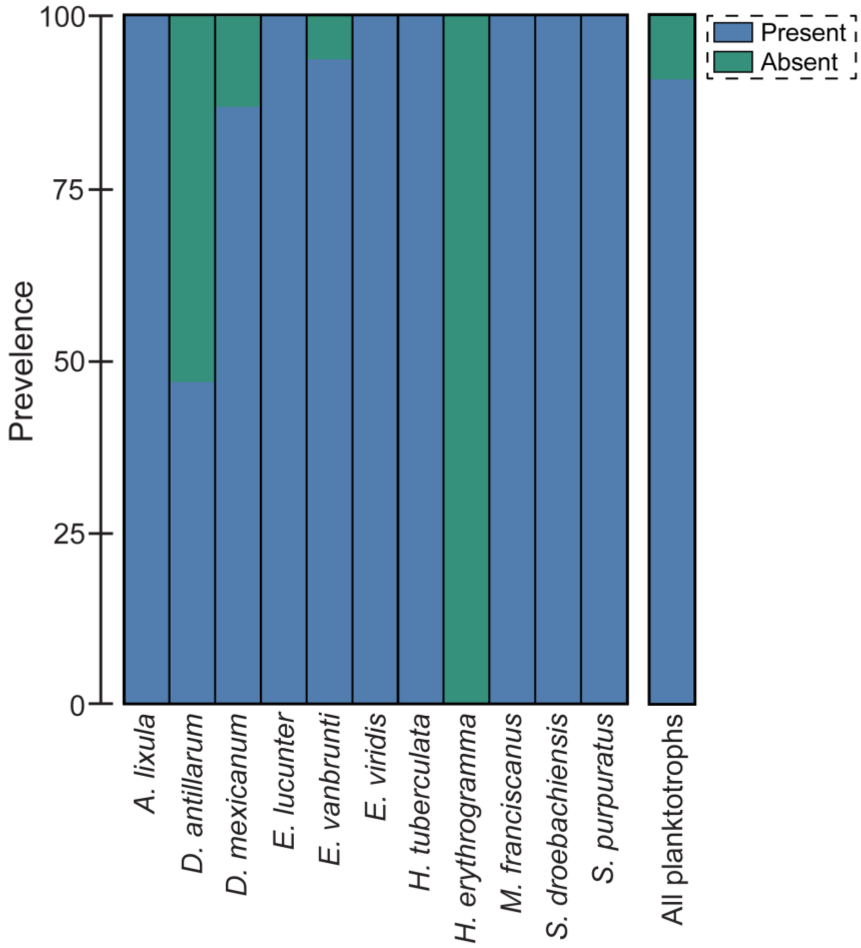


**Fig. S3:** Prevalence of plastid ASVs in sea urchin eggs. Plastid ASVs were present in the eggs of all but one sea urchin species, which was the only sea urchin that develops and undergoes metamorphosis without the requirement of external nutrients through feeding. The other species develop via feeding larvae, of which plastid DNA was present in 91% of these samples. Note: the two *Eucidaris* species in Fig. S2 were not included in the rest of this meta-analysis because the single sample for each is insufficient for a full comparison.

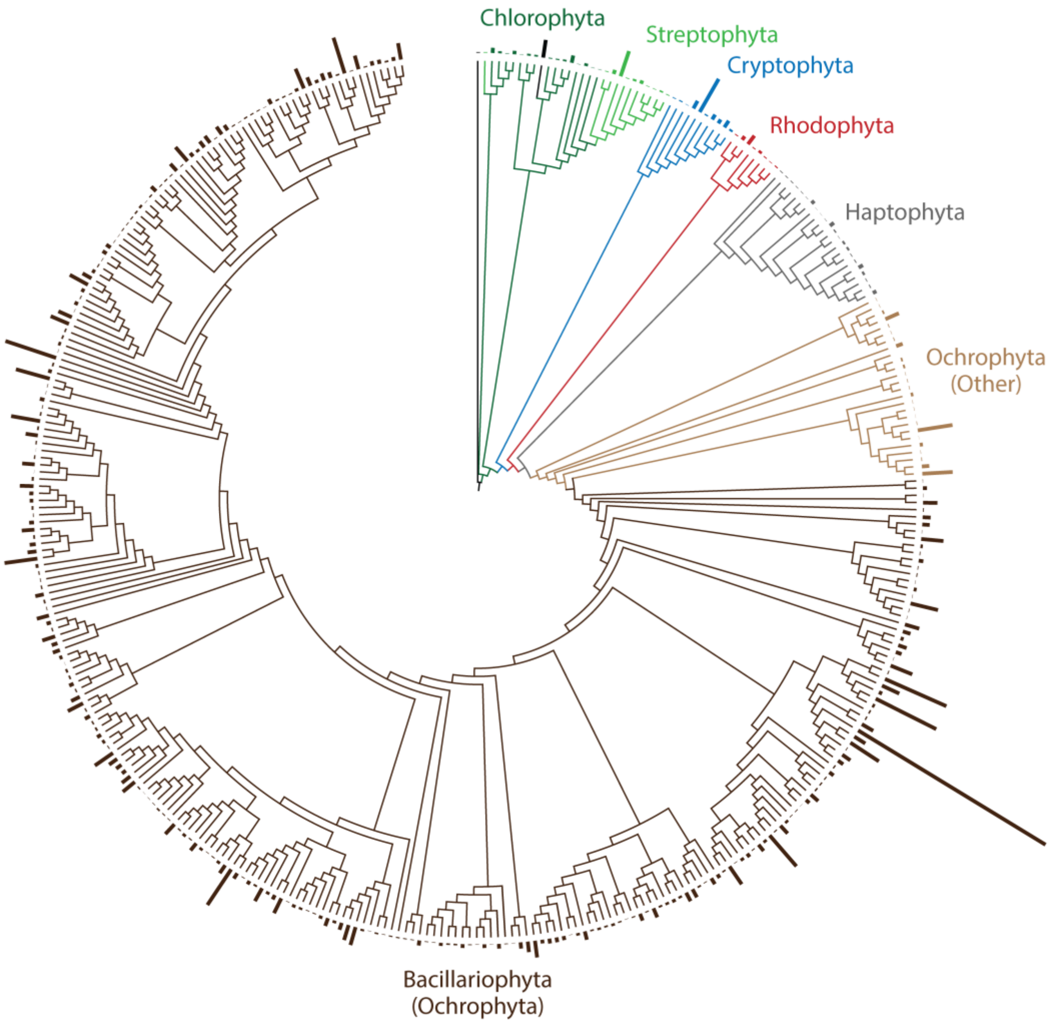


**Fig. S4**: Plastid ASVs are predominately diatoms. Sea urchins collectively provide hundreds of plastid ASVs to their eggs. Apicomplexa (black leaf in the center) and Discoba (black leaf to the right) were represented by a single ASVs, Chlorophyta (dark green), Cryptophyta (blue), Haptophyta (gray), Rhodophyta (red), and Streptophyta (light green) were represented by 10 to 22 ASVs each, and the Ochrophyta (brown) were represented by 308 ASVs. The major of Ochrophyta ASVs were derived from the Bacillariophyta (*i.e.*, diatoms; dark brown). Bars on the outer ring represents the relative abudance of each ASV.

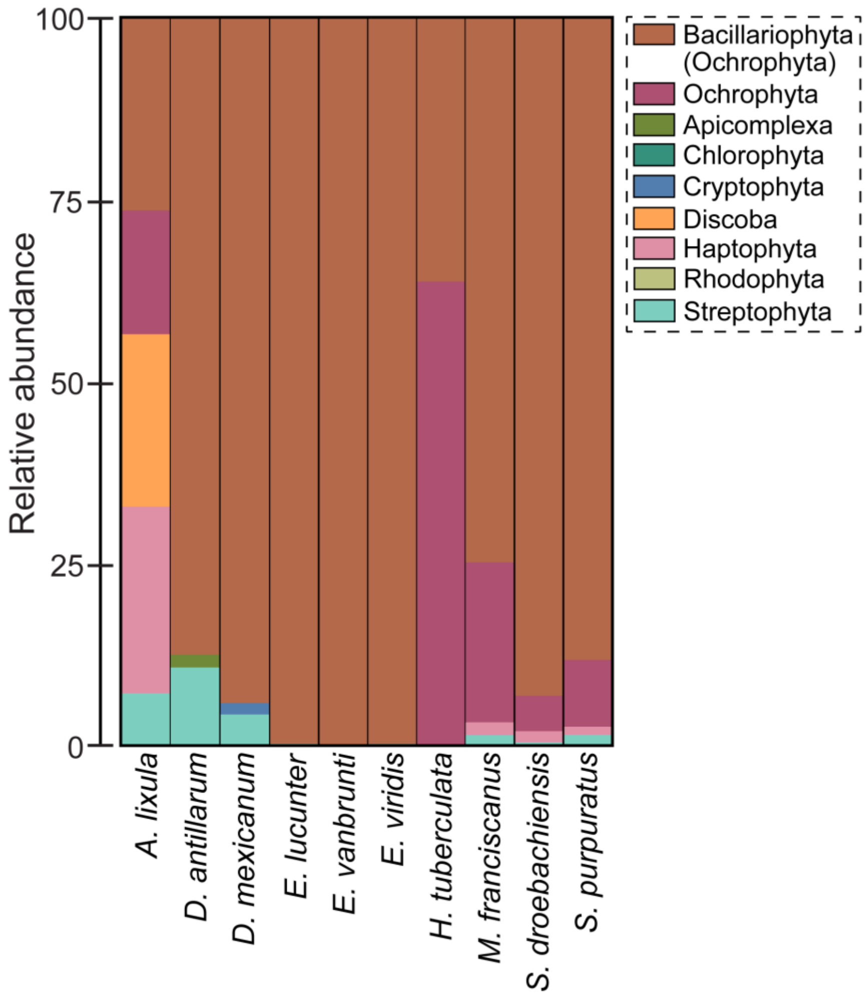


**Fig. S5:** Taxonomic profile of plastid ASVs. Taxonomic profile of the plastid ASVs from photosynthetic eukaryotes in sea urchin eggs. All groups represent eukaryotic phyla except the Bacillariophyta (diatoms), which are presented at the class-level and is part of the Ochrophtya.

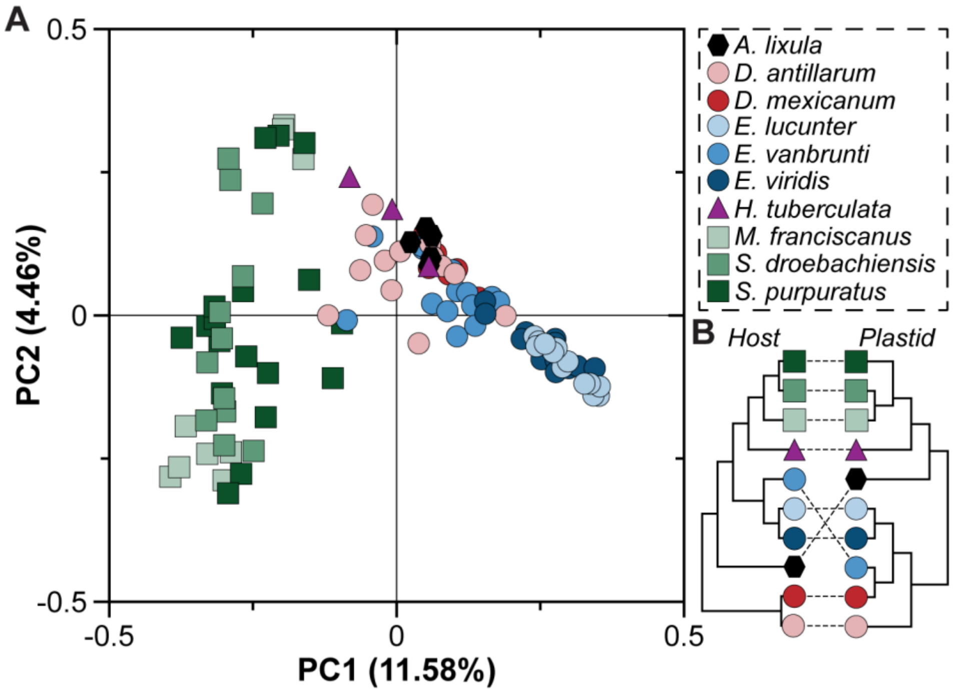


**Fig. S6:** Species and location-specificity in plastid ASVs. (A) Principal coordinate analysis depicting community relatedness (Jaccard) of the plastid ASVs in sea urchin eggs. Color and shape denote species and geographic locations, respectfully. Square, circle, triangle, and hexagon represent Friday Harbor (WA, USA), Panama (Panama City and Colón), Sydney (Australia), and Tenerife (Canary Islands, Spain), respectfully. (B) Topology of the host gene tree (using COI) is congruent with a dendrogram for these plastid ASVs based on species, even though they do not fully mirror each other.

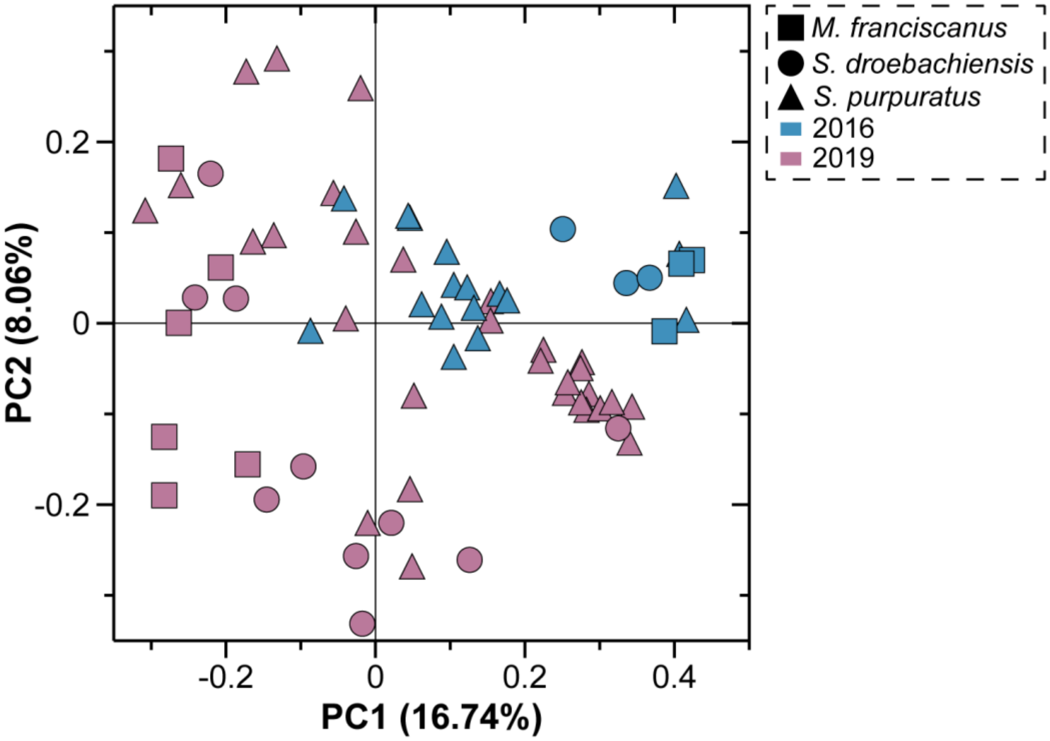


**Fig. S7:** Temporal differences in plastid ASVs. Principal coordinate analysis depicting community relatedness (Jaccard) of plastid ASVs that are associated with the eggs of the confamilials *Strongylocentrotus purpuratus* (triangle), *Mesocentrotus franciscanus* (square), and *S. droebachiensis* (circle) that were collected in 2016 (blue) and 2019 (pink) in Friday Harbor (WA, USA).

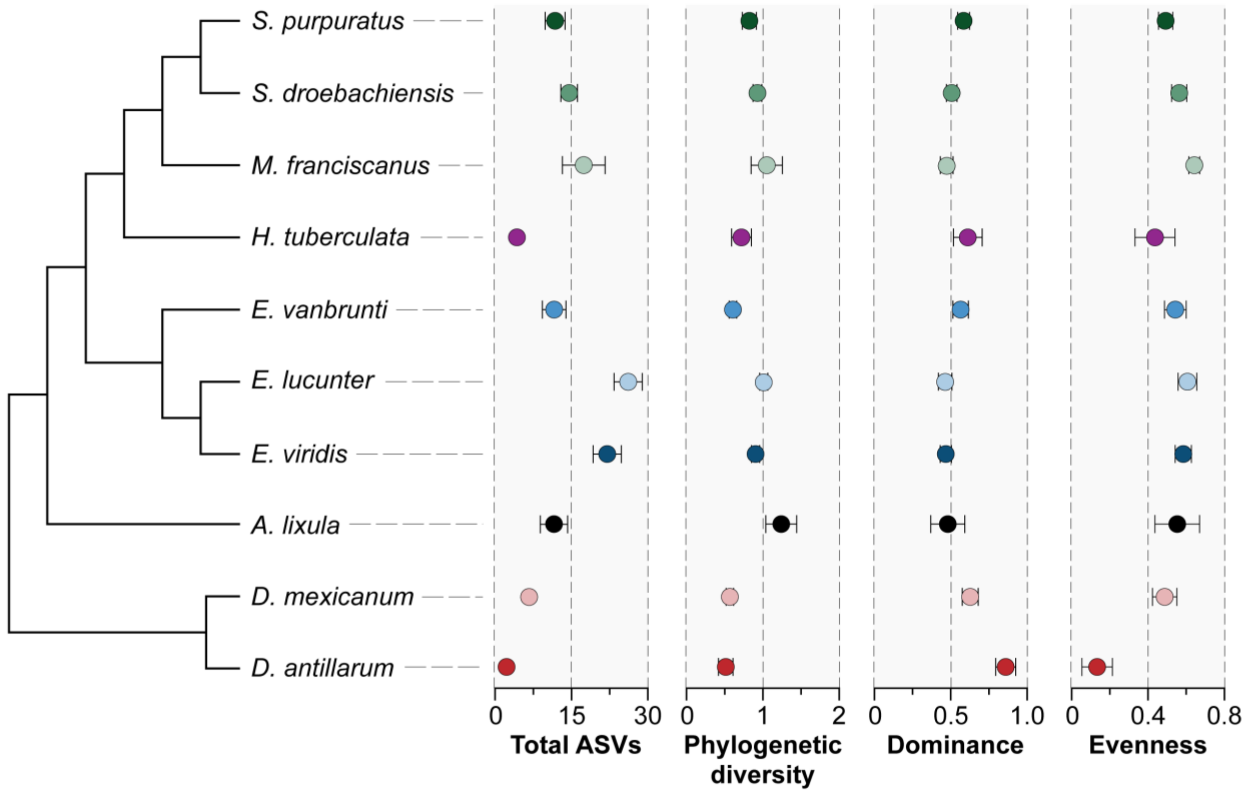


**Fig. S8:** Plastid diversity in sea urchin eggs. This was estimated by total ASVs, Faith's phylogenetic diversity, McIntosh dominance, and McIntosh evenness. Each circle species average ± standard error.

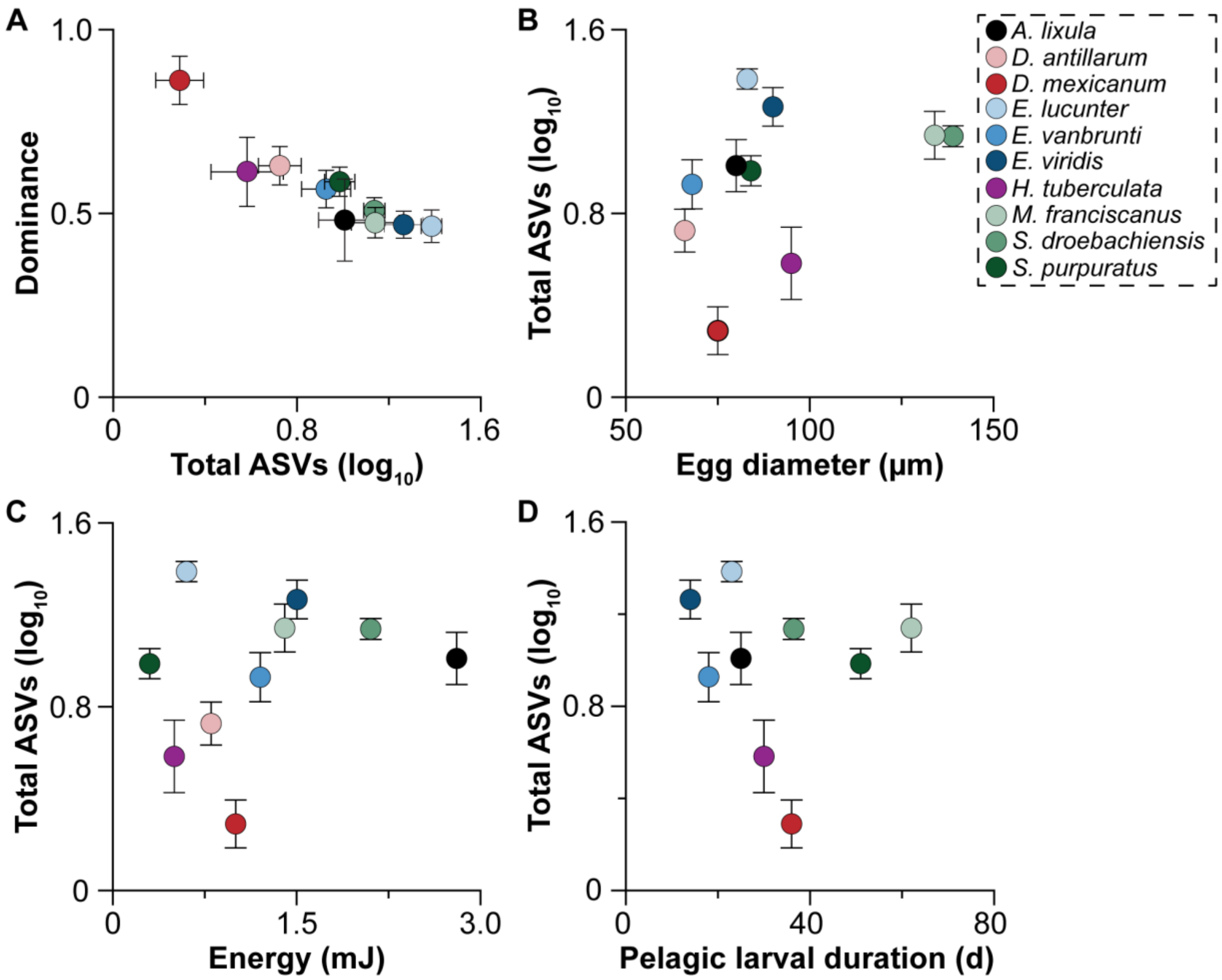


**Fig. S9:** Egg size is proportional to plastid ASV richness. (A) ASV diversity and dominance within the collection of plastid ASVs provided to sea urchin eggs were negatively correlated. (B) Egg diameter and the richness of plastid ASVs that were provided to sea urchin eggs are positively correlated. (C, D) No trade-off was observed between the energetic content (mJ; linear regression: $F_{1,109} = 1.36$, $p = 0.246$, $R^2 = 0.012$) of sea urchin eggs or pelagic larval duration (days; linear regression: $F_{1,109} = 0.380$, $p = 0.539$, $R^2 = 0.003$) and the diversity of plastid ASVs that were provided to sea urchin eggs. Each circle represent a species average with their corresponding standard error bar.

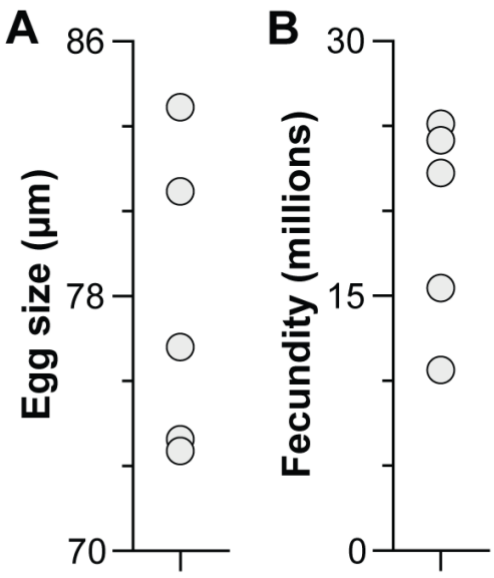


**Fig. S10:** Egg size and fecundity. Average egg size (diameter; µm) (A) and fecundity (B) for five individuals of the sea urchin *Arbacia lixula*.

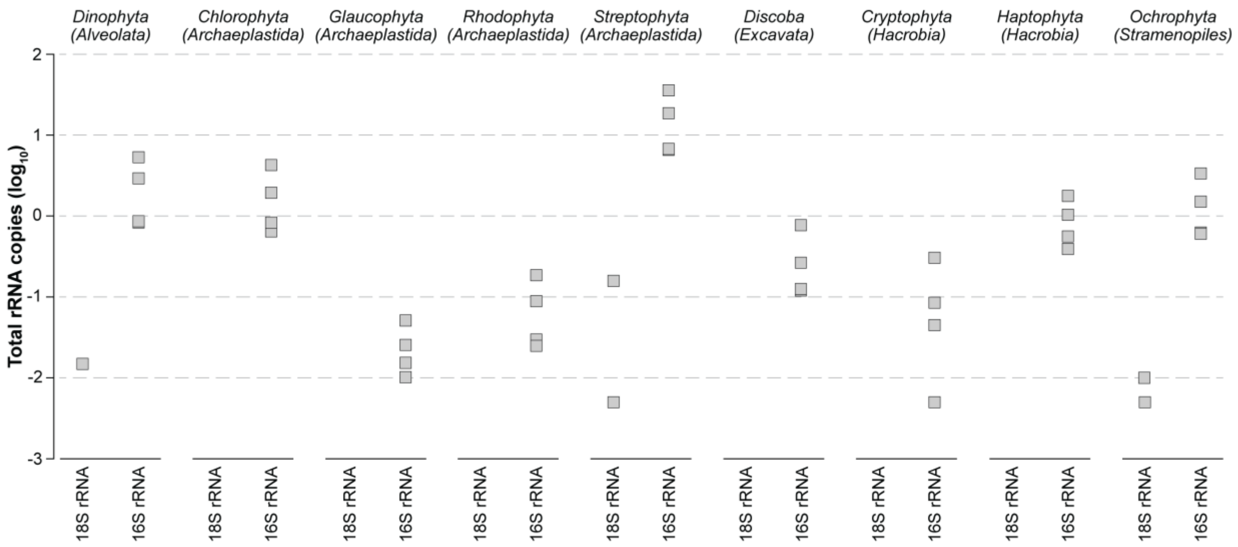


**Fig. S11:** Lack of a nuclear gene for most photosynthetic eukaryotes. A nuclear gene marker (18S rRNA gene) was disproportionately low in abundance or not present in embryos of the sea urchin *Arbacia lixula* compared to a plastid gene marker (16S rRNA gene) for the Alveolata (Dinophyta), Archaeplastida (Chlorophyta, Glaucophyta, Rhodophyta, Streptophyta), Excavata (Discoba), Hacrobia (Cryptophyta, Haptophyta), and Stramenopiles (Ochrophyta). This is a group-by-group display of the rRNA counts presented in Fig. 1C and the raw data presented in Table S8.

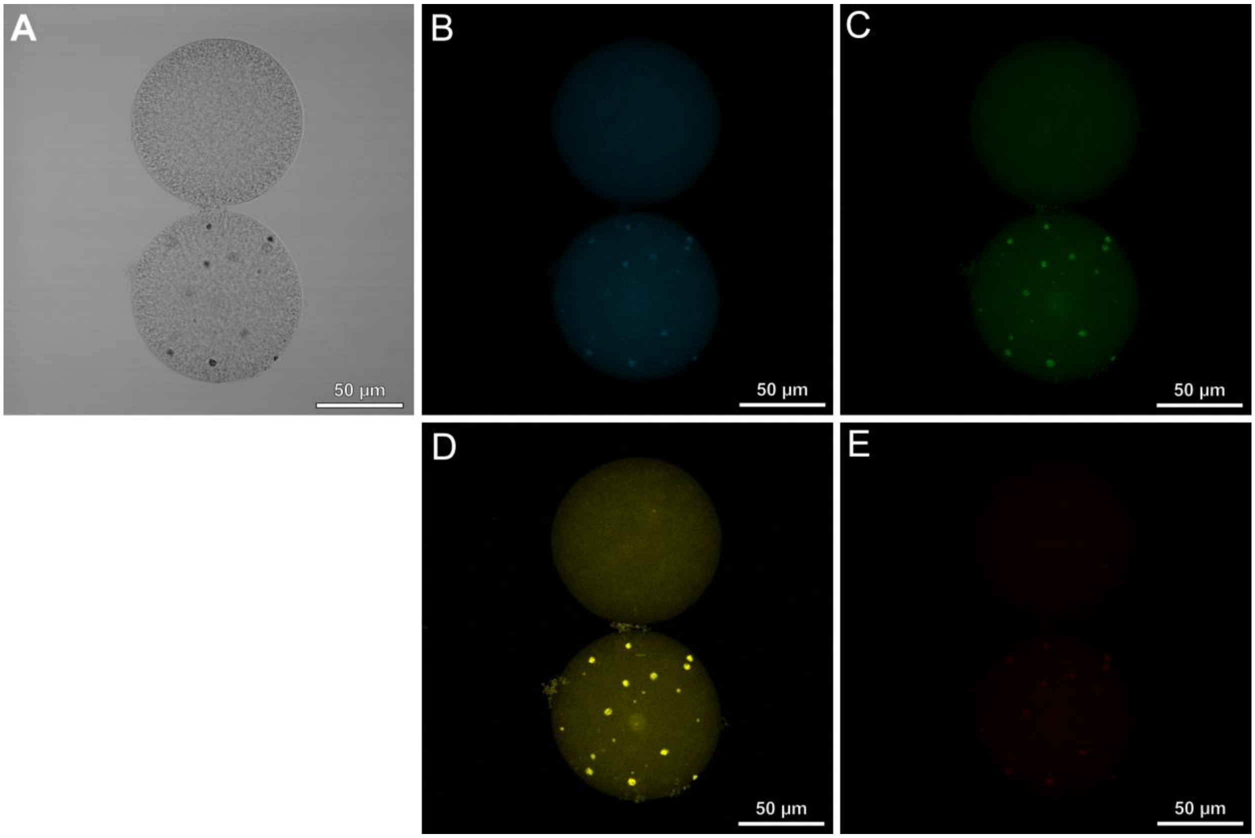


**Fig. S12:** Cytoplasmic autofluorescent structure in eggs. Light micrographs of eggs from the sea urchin *Arbacia lixula* without (top) and with (bottom) cytoplasmic autofluorescent particles, as displayed by transmitted light (A) as well as at an excitation of 405 nm (B), 488 nm (C), 561 nm (D), and 640 nm (E). The light micrograph is a single optical slice, while the florescence micrograph is a maximum intensity projection of a Z-stack. Note: A and D are identical to Fig. 2A and 2B, respectfully. These are displayed here again to allow for direct comparison across wavelengths.

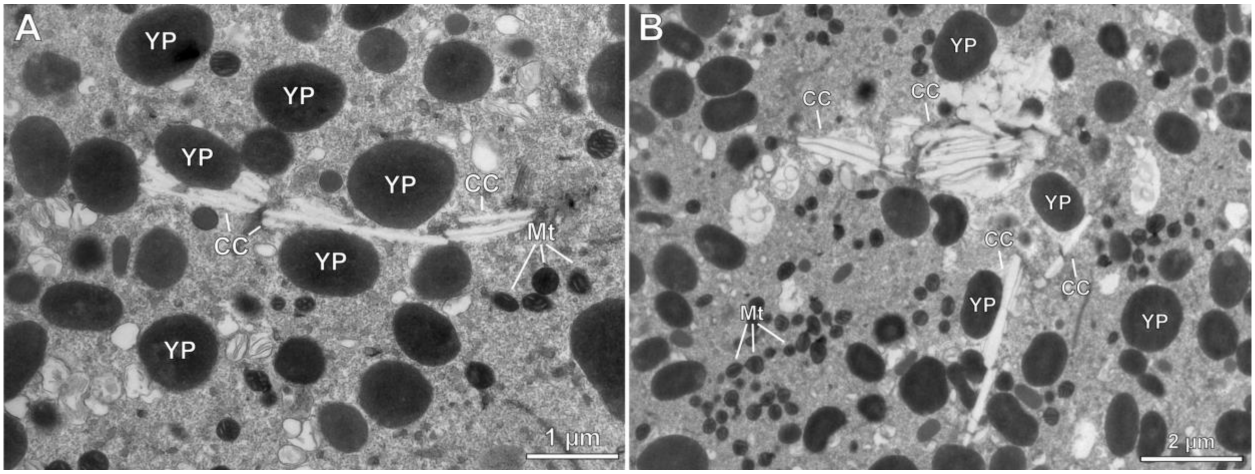


**Fig. S13:** Diversity of this crystal structure. Transmission electron micrographs of eggs from the sea urchin *Arbacia lixula* with what we presume are large tetragonal carotenoid crystals (CC). Chromoplast-specific starch granules and plastoglobules could not be distinguished within *A. lixula* eggs. Also in view are yolk platelets (YP) and mitochondria (Mt) that are presumed to be maternal.

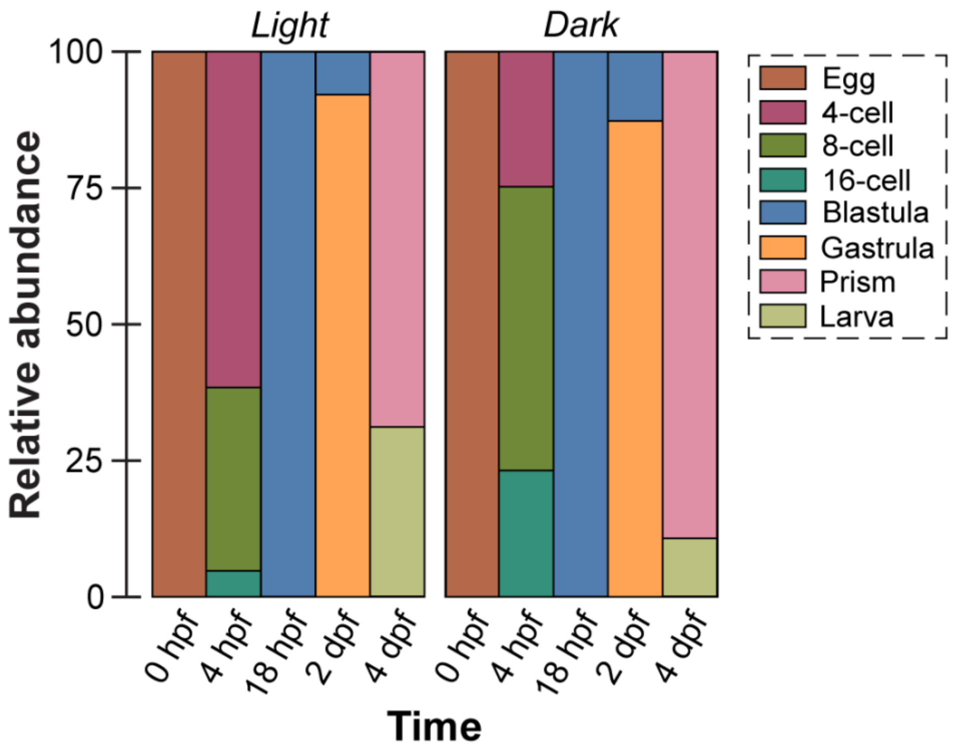


**Fig. S14:** Summary of developmental profiles. The developmental stages for offspring of the sea urchin *Arbacia lixula* were recorded at several intervals while being cultured from fertilization through four days post-fertilization in light (*i.e.*, with the light-dependent activity of chromoplast-derived components) and dark (*i.e.*, without the light-dependent activity of chromoplast-derived components).

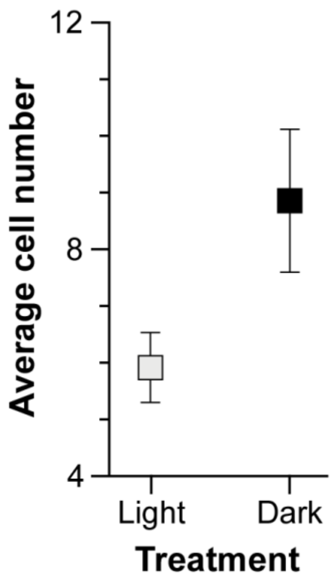


**Fig. S15:** Increase in cell division. Offspring develop quicker in dark (*i.e.*, without the light-dependent activity of chromoplast-derived components) during the first four cell divisions (*i.e.*, from fertilization to the 16-cell stage), as compared to their siblings in light (*i.e.*, with the light-dependent activity of chromoplast-derived components). All values are average ± standard error.

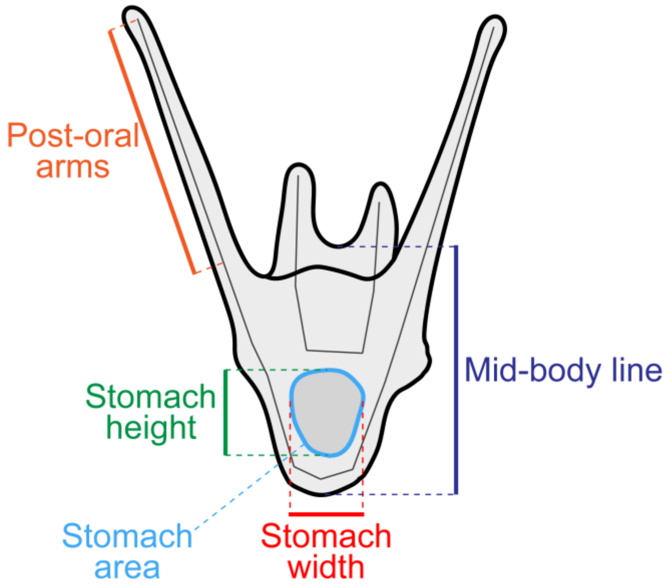


**Fig. S16:** Cartoon larva. Schematic of the morphological features that were measured to assess whether the two-arm larvae (four days post-fertilization) of the sea urchin *Arbacia lixula* exhibited morphological plasticity.

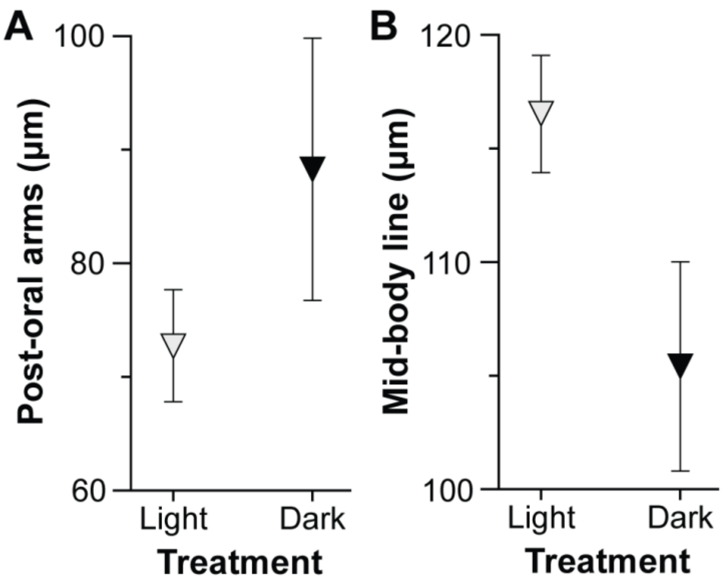


**Fig. S17:** Light-induced developmental plasticity for some morphological features. The length of the post-oral arms (A) and the mid-body line (B) were similar between treatments for larvae of the sea urchin *Arbacia lixula* that were cultured in light (*i.e.*, with the light-dependent activity of chromoplast-derived components), as compared to their siblings in dark (*i.e.*, without the light-dependent activity of chromoplast-derived components). All values are average ± standard error.

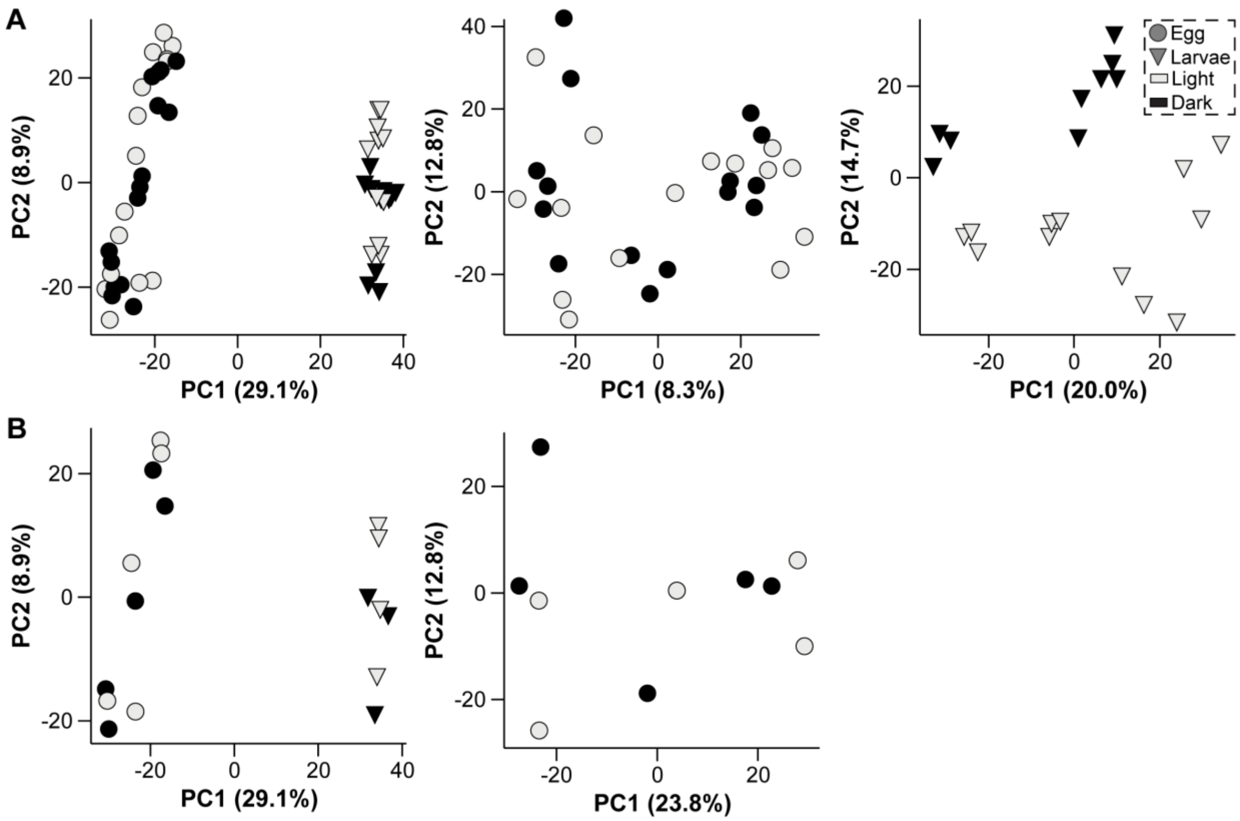


**Fig. S18:** Light-induced metabolic divergence during development. Eggs of the sea urchin *Arbacia lixula* that were placed in light (*i.e.*, with the light-dependent activity of chromoplast-derived components) and dark (*i.e.*, without the light-dependent activity of chromoplast-derived components) had similar metabolomes (left and center). A metabolome-wide shift occurred during development (left) and based on the presence of light for larvae (right). These PCA plots are displayed with each technical replicate (A) as well as the median of those technical replicates (B). Larvae with the median of their technical replicates is Fig. 3D.

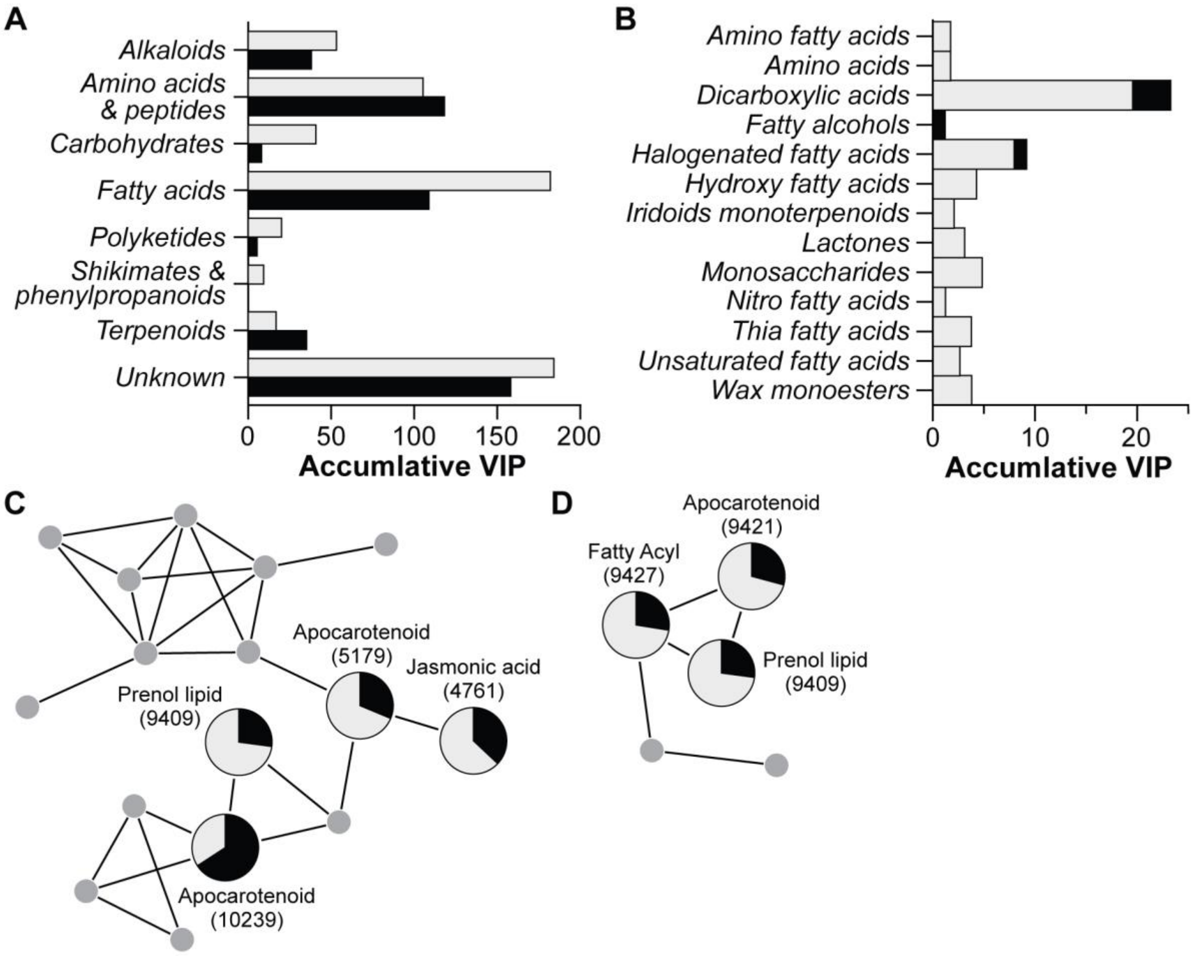


**Fig. S19:** Differentially abundant metabolites and apocarotenoid modules. (A) Larvae cultured in light and dark exhibit organism-wide differences in their metabolism. (B) This was predominately driven by a shift in fatty acid metabolism, including dicarboxylic acids as well as several other complementary fatty acids that were mainly differentially abundant in light. (C) Metabolite 9409 (a prenol lipid) was part of a metabolic module that had three other known metabolites that were found in the offspring of the sea urchin *Arbacia lixula*. This module included two uncharacterized apocarotenoids and jasmonic acid, all of which are phytohormones. (D) Metabolite 9421 was part of a metabolic module that included a prenol lipid and fatty acyl.

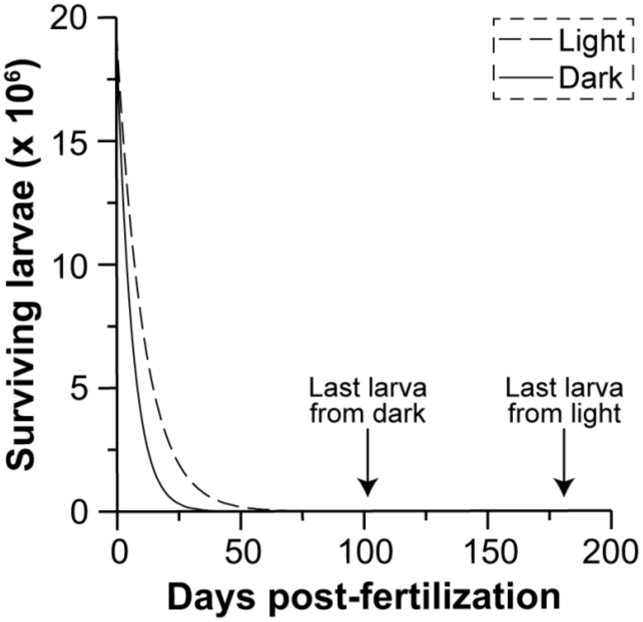


**Fig. S20:** Enhanced dispersal potential. Planktonic duration was estimated for development in light (*i.e.*, with the light-dependent activity of chromoplast components) and dark (*i.e.*, without the light-dependent activity of chromoplast components) using the fecundity of the sea urchin *Arbacia lixula* and the instantaneous rate model for the natural mortality of marine invertebrate larvae. It is estimated to take 181 and 102 days in light and dark, respectfully, for the last larva from an entire clutch of an individual *A. lixula* to remain.

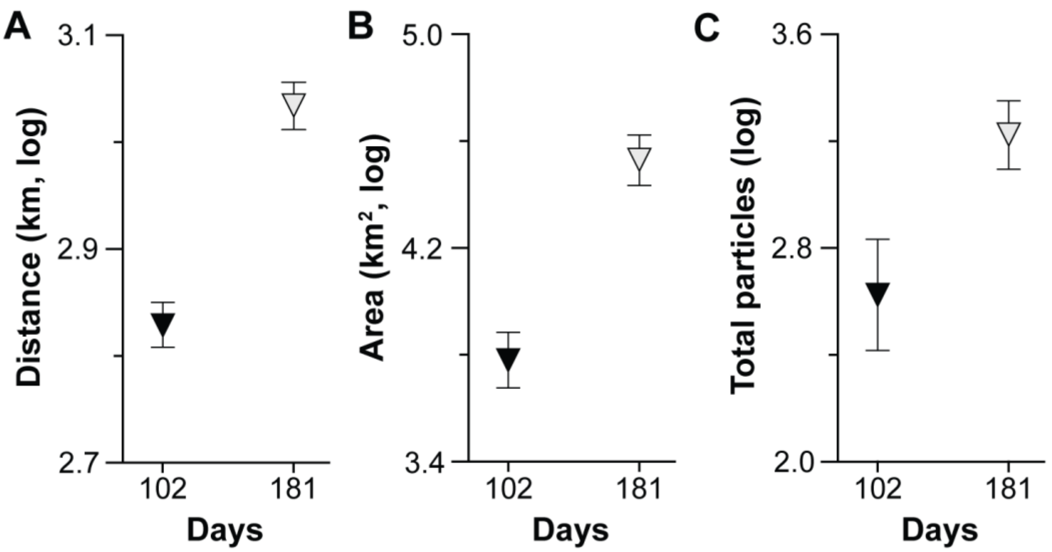


**Fig. S21:** Enhanced dispersal. Offspring cultured in light (*i.e.*, with the light-dependent activity of chromoplast components) enhances the total dispersal distance (A), geographical area that is settled on (B), and abundance of particles reaching a habitat to settle (C), as compared to their siblings in dark (*i.e.*, without the light-dependent activity of chromoplast components). These numerical values were extracted from Fig. 4A, of which are derived from the distribution of particles on the African shelf after 102 (*i.e.*, without the light-dependent activity of chromoplast components) and 181 (*i.e.*, with the light-dependent activity of chromoplast components) days of being released from Tenerife (Canary Islands, Spain). All data were log-transformed for normality. Each dot represents the 10-year average ± standard error.

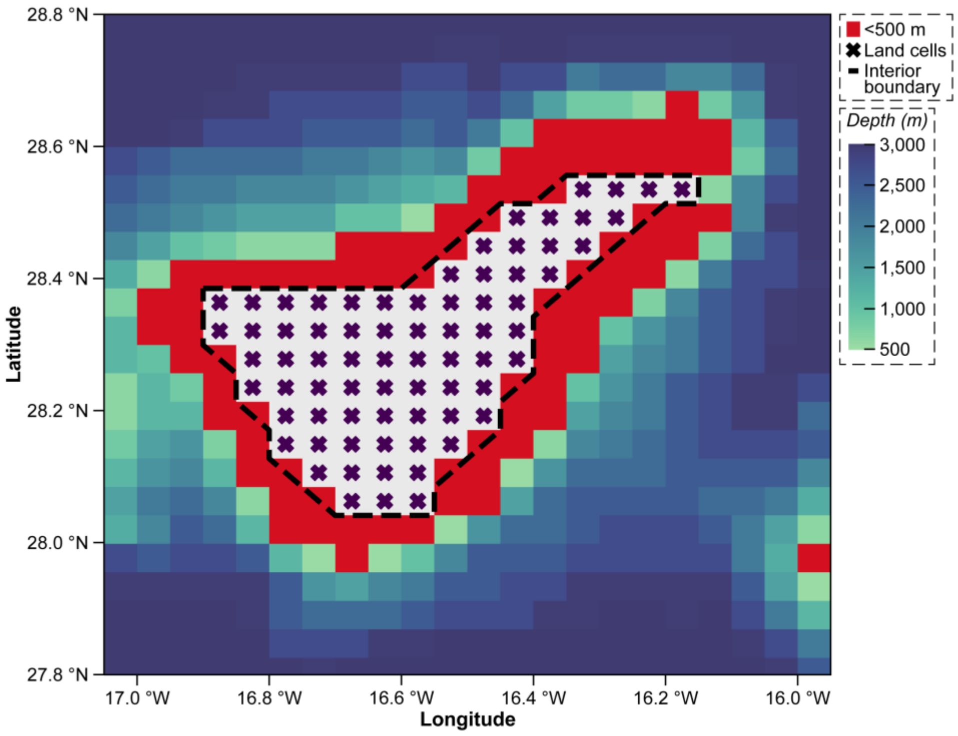


**Fig. S22:** Location for particle release. Location of particle release from Tenerife (Canary Islands, Spain) at the grid resolution of the VIKING20X model. Particles were released from grid cells that were <500 m (red squares) adjacent to Tenerife.

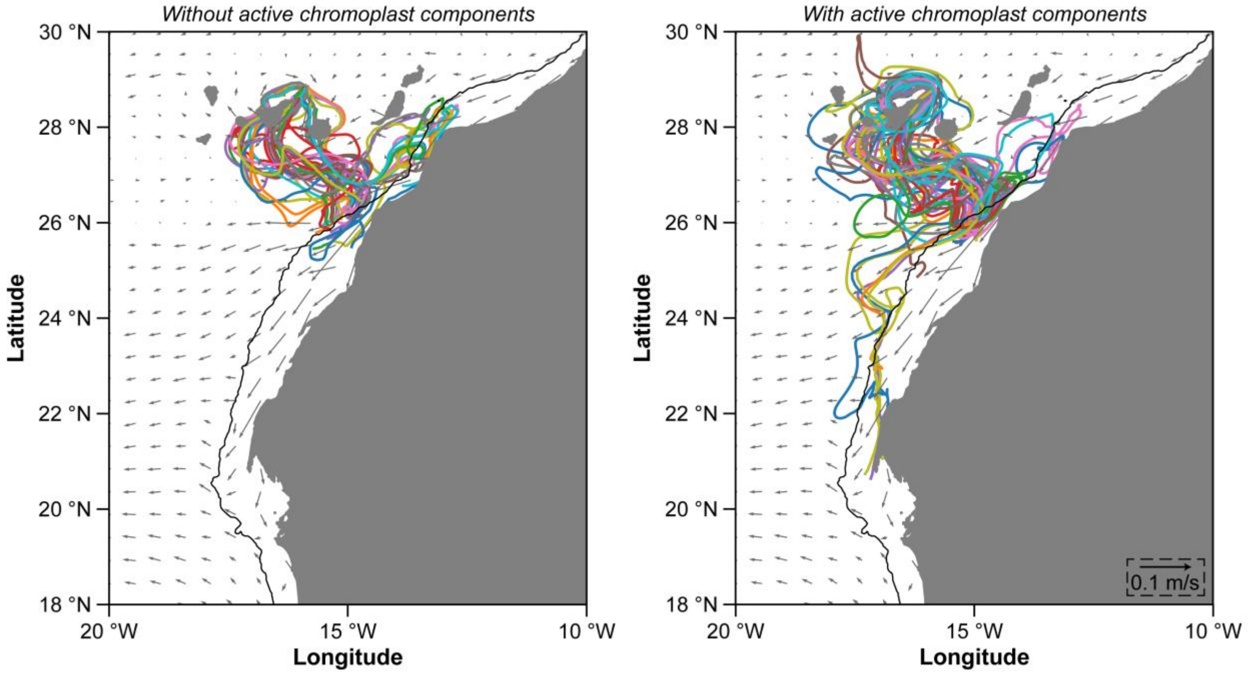


**Fig. S23:** Particle trajectories. Example particle trajectories (n = 50) from Tenerife (Canary Islands, Spain) to the African shelf for dispersal durations of 102 (*i.e.*, dark, without the light-dependent activity of chromoplast components; left) and 181 (*i.e.*, light, with the light-dependent activity of chromoplast components; right) days. Grey arrows represent the 10-year mean (2007-2016) velocity for the release depth (0-30 m) from the VIKING20X model. Every tenth arrow is shown. Black contours mark the 500 m isobath around Africa.

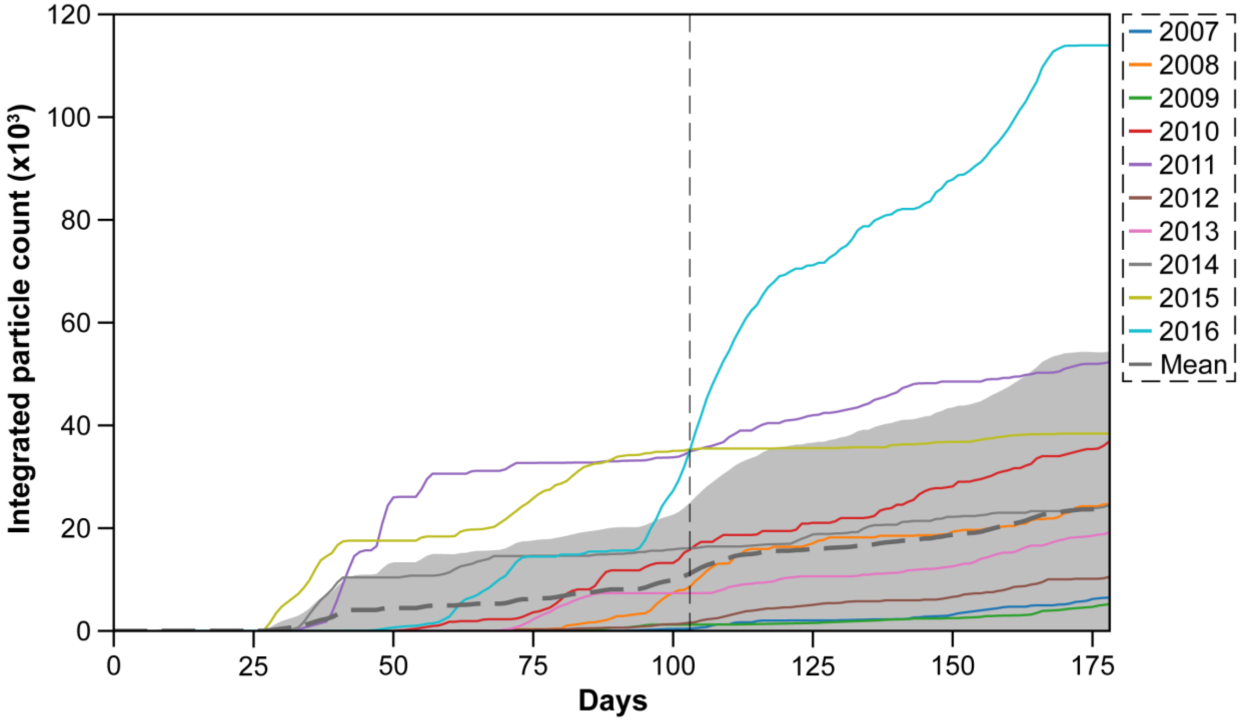


**Fig. S24:** Annual variation. Annual variation in particles reaching the African Shelf, as based on the VIKING20X model. The dashed gray line represents the mean for years 2007 to 2016 and the shaded areas mark one standard deviation. The vertical line indicates 102 days after release (*i.e.*, dark, without the light-dependent activity of chromoplast components).

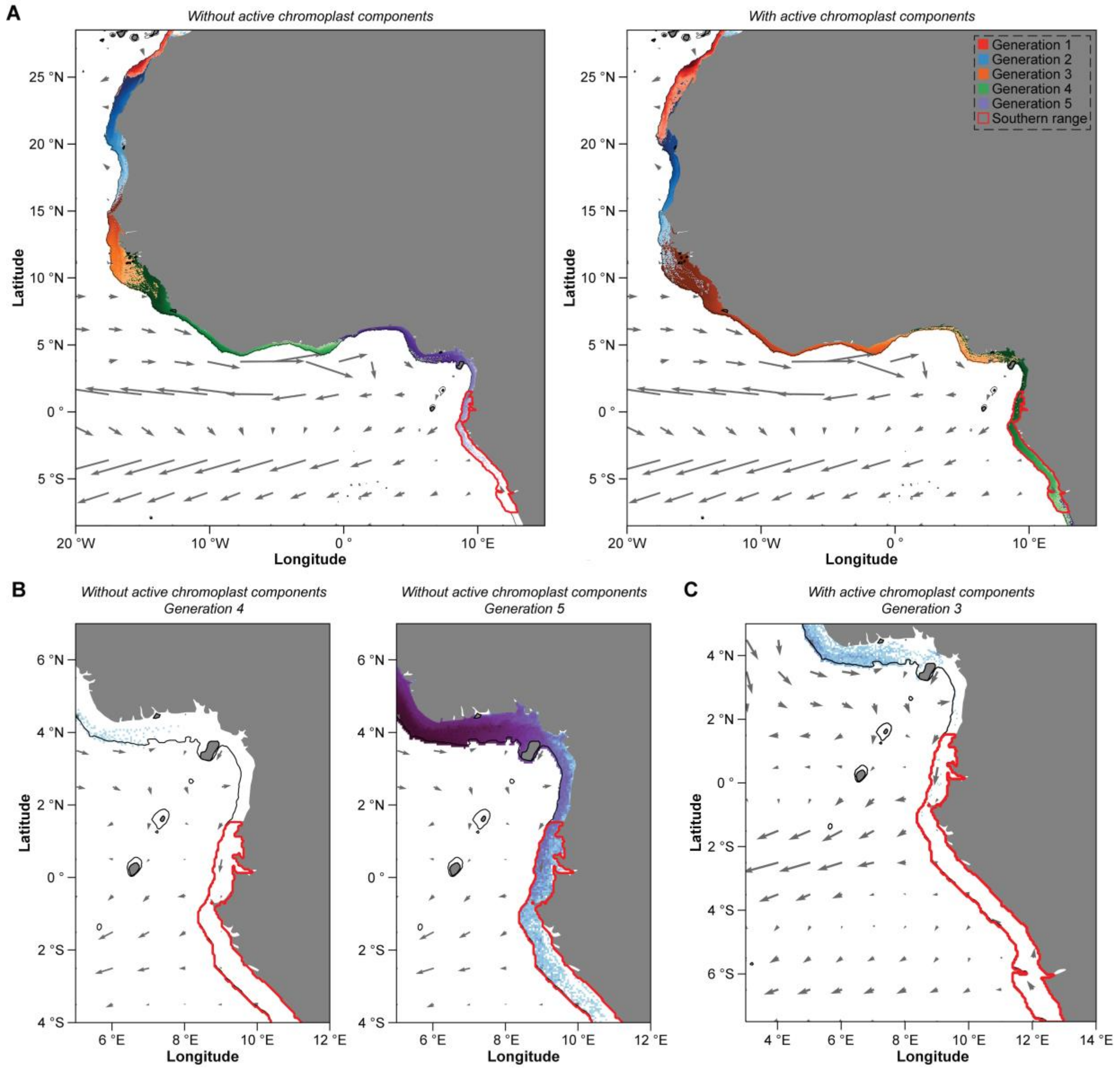


**Fig. S25:** Offspring spread more quickly. Modeled distribution of particles on the African shelf over multiple generations of dispersal. (A) Initial particles (*i.e.*, Generation 0) were released from Tenerife (Canary Islands, Spain), and were provided 102 (*i.e.*, dark, without the light-dependent activity of chromoplast components; left) or 181 (*i.e.*, light, with the light-dependent activity of chromoplast components; right) days to disperse. A second dispersal event of 102 or 181 days then initiated from their respected distributions. This was repeated until particles in each treatment reached the southernmost area of the range for the sea urchin *Arbacia lixula* (*i.e.*, the red area). (B) Offspring without the light-dependent activity of chromoplast components were close to reaching this on the Generation 4 (left) and did reach it on Generation 5, while those with the light-dependent activity of chromoplast components reached this area on Generation 3 (C). Grey arrows represent the 10-year mean velocity for the release depth (0-30 m) from the VIKING20X. Every tenth arrow is shown. Black contours around the African shelf mark the 500 m isobath.

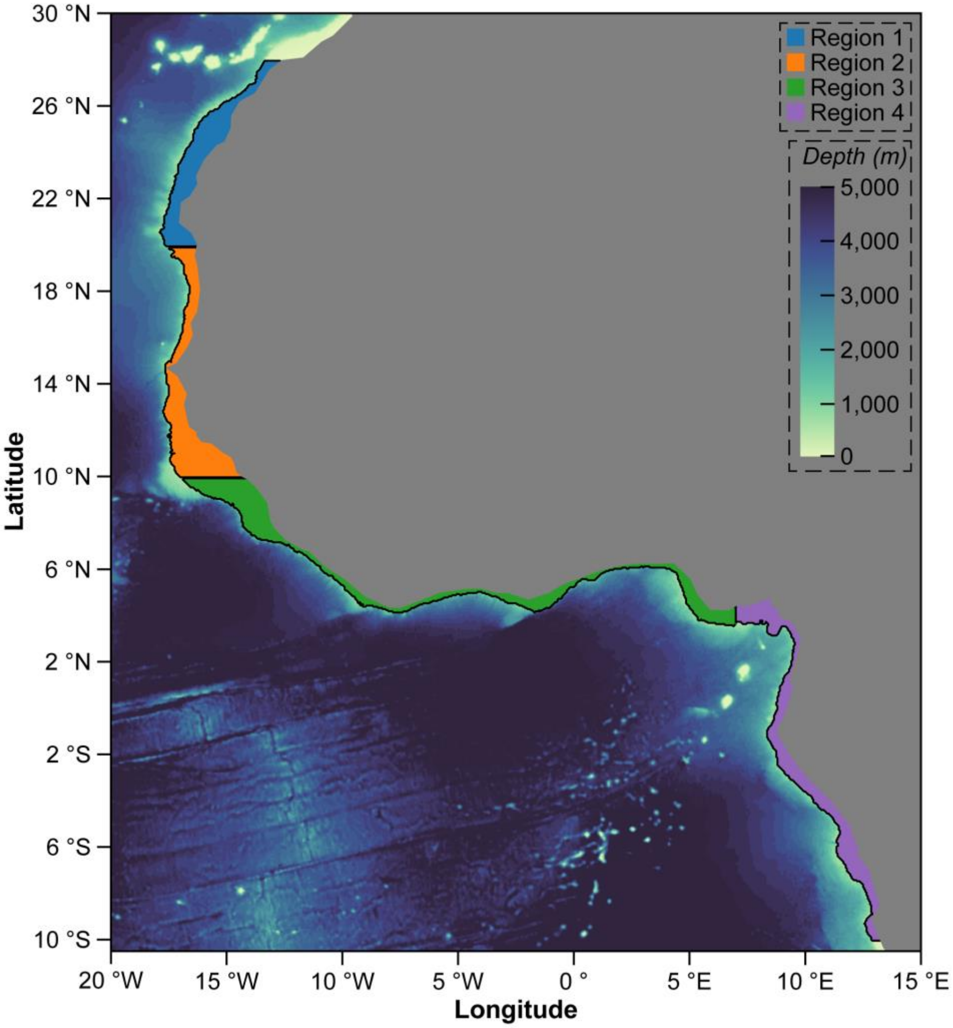


**Fig. S26:** Zones of particle release off the coast of Africa. Particles were released from each of the four regions off the coast of Africa to determine if offspring with and/or without the light-dependent activity of chromoplast components could disperse across the Atlantic Ocean. Black contours mark the 500 m isobath around the African Coast.

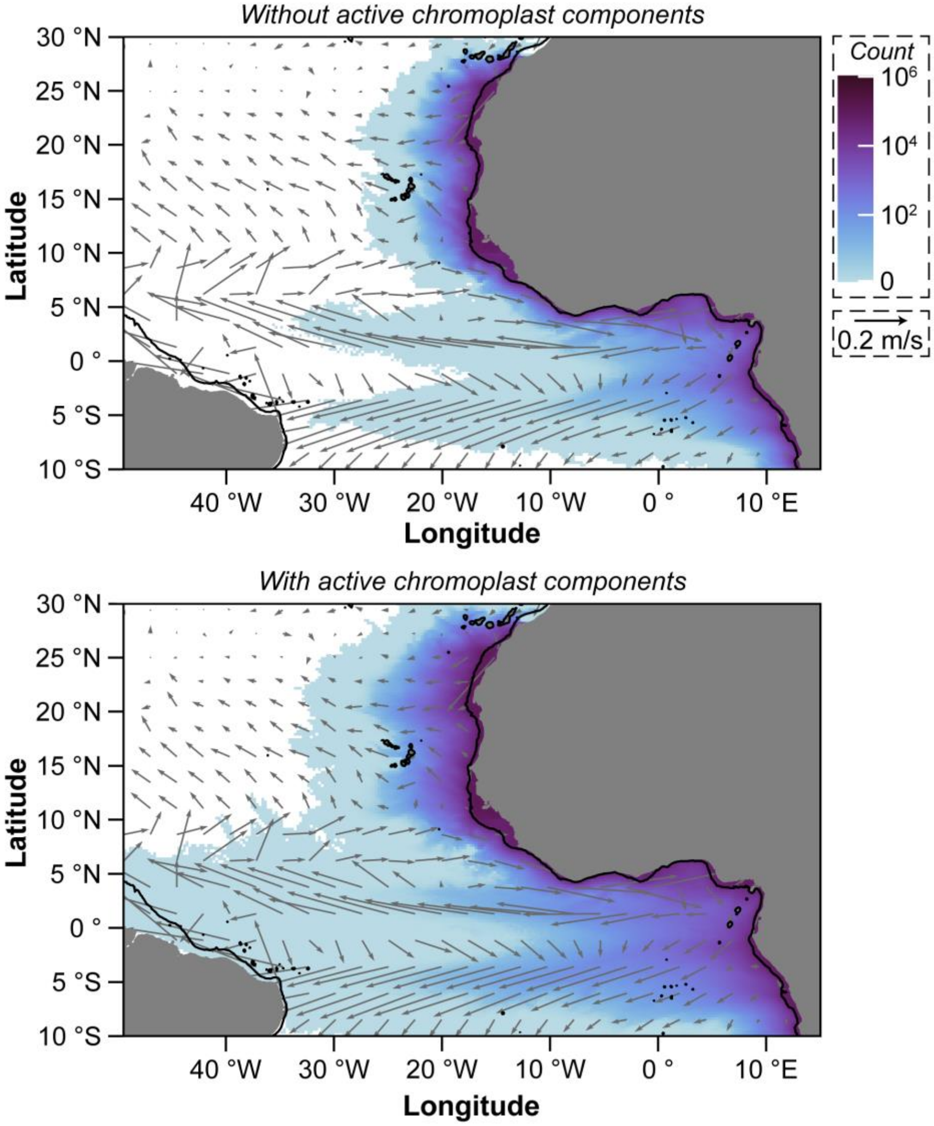


**Fig. S27:** Enabling trans-Atlantic dispersal. Distribution of particles in the Atlantic Ocean along the coast of Africa after 102 (*i.e.*, dark, without the light-dependent activity of chromoplast components; left) and 181 (*i.e.*, light, with the light-dependent activity of chromoplast components; right) days of being released. Grey arrows represent the 10-year mean velocity for the release depth (0-30 m) from VIKING20X. Every 50$^{th}$ arrow is shown. Black contours mark the 500 m isobath around Africa.

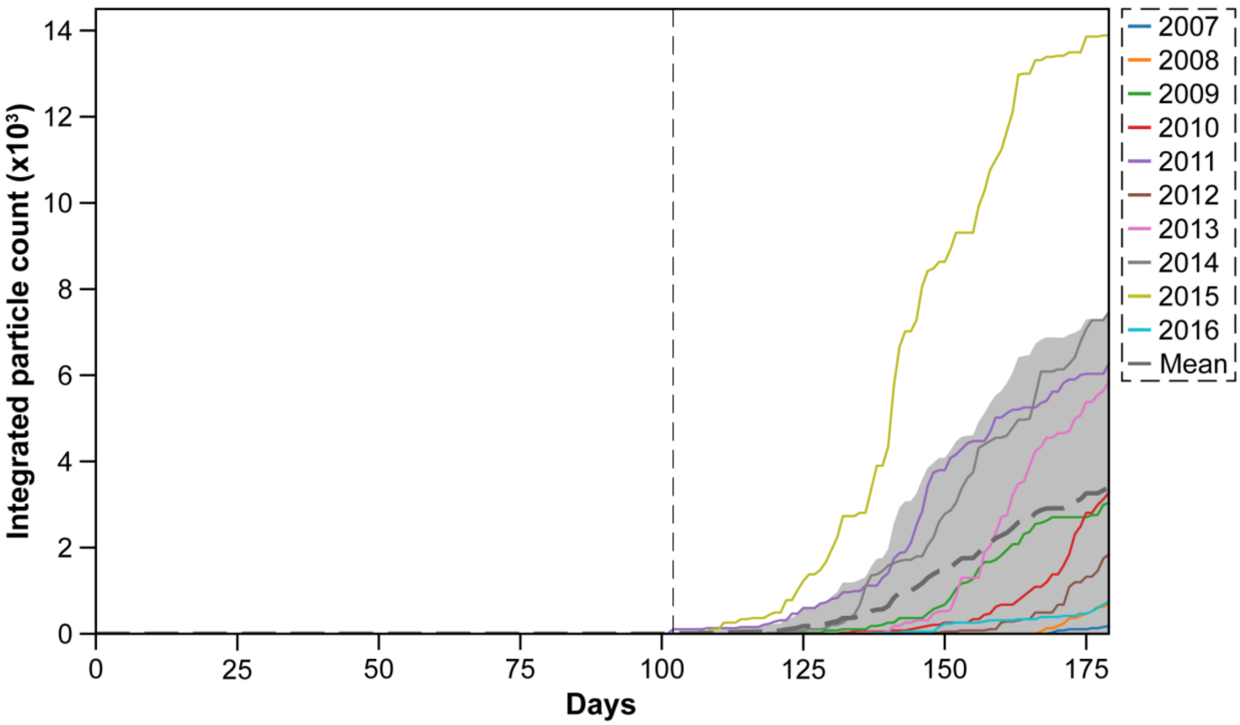


**Fig. S28:** Annual variation. Annual variation in particles that disperse from the African Shelf to Brazil, as based on the VIKING20X model. The dashed gray line represents the mean for years 2007 to 2016 and the shaded areas mark one standard deviation. The vertical line indicates 102 days after release (*i.e.*, dark, without the light-dependent activity of chromoplast components).

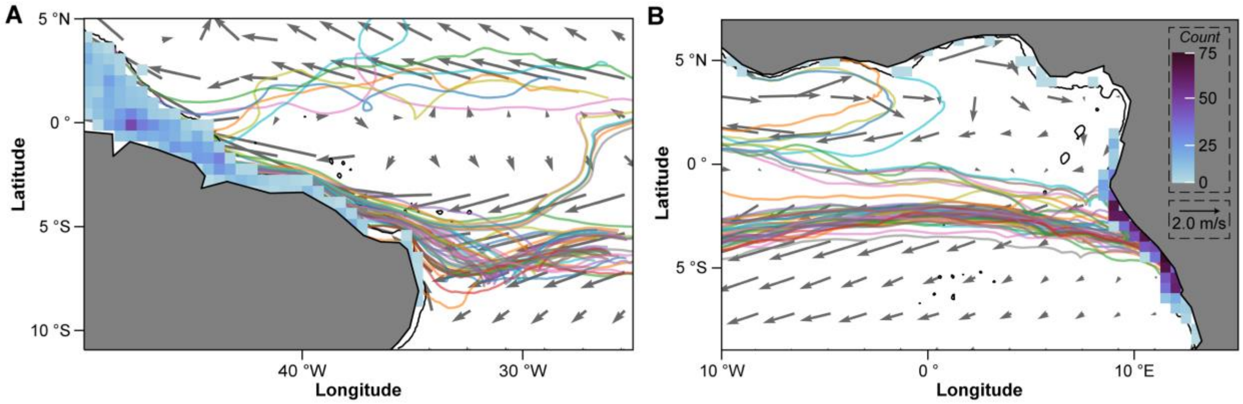


**Fig. S29:** Paths across the Atlantic Ocean. Distribution and abundance of particles that reach the coast of Brazil (A) and that depart from the coast of Africa (B) after 181 days of being released (*i.e.*, light, with the light-dependent activity of chromoplast components). This is a zoomed-in version of Fig. 4B. Grey arrows represent the 10-year mean velocity for the release depth (0-30 m) from VIKING20X. Every 50$^{th}$ arrow is shown. Black contours mark the 500 m isobath around Africa.

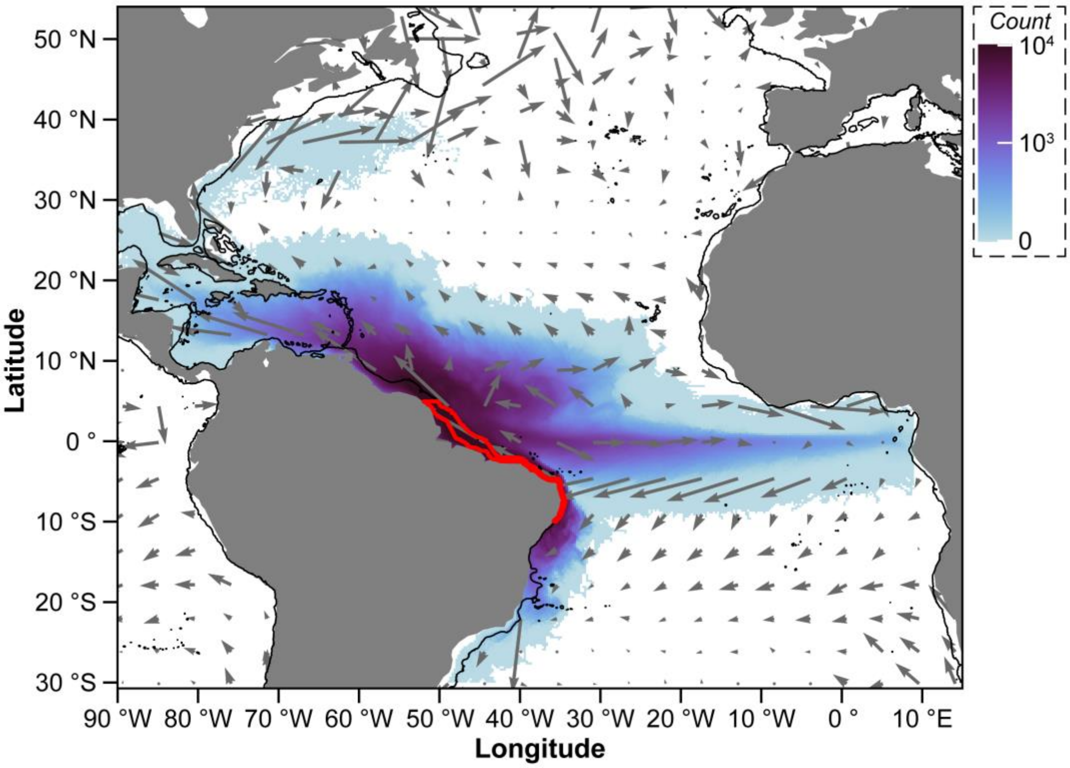


**Fig. S30:** Dispersal from Brazil. Distribution of particles in the Atlantic Ocean that were released along the northern coast of Brazil (red) after 181 (*i.e.*, light, with the light-dependent activity of chromoplast components) days of being released. Grey arrows represent the 10-year mean velocity for the release depth (0-30 m) from VIKING20X. Every $100^{th}$ arrow is shown. Black contours represent 500 m isobath.

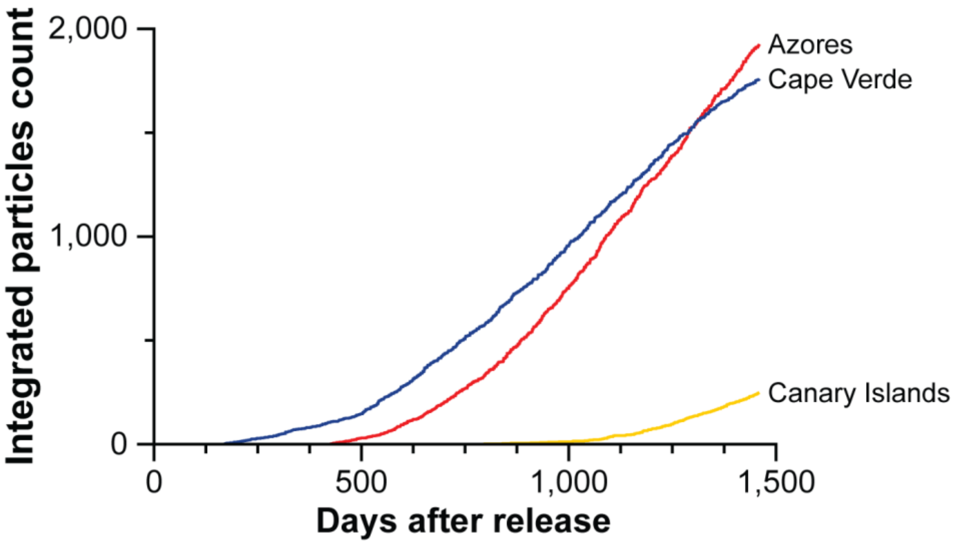


**Fig. S31:** Unable to reach Macaronesia. Duration required for particles to disperse from the northern coast of Brazil to Macaronesia. A particle was first detected in Cape Verde on day 170, the Azores on day 366, and the Canary Islands on day 783. This is based on the VIKING20X model and is the accumulation of annual releases from years 2007 to 2016.

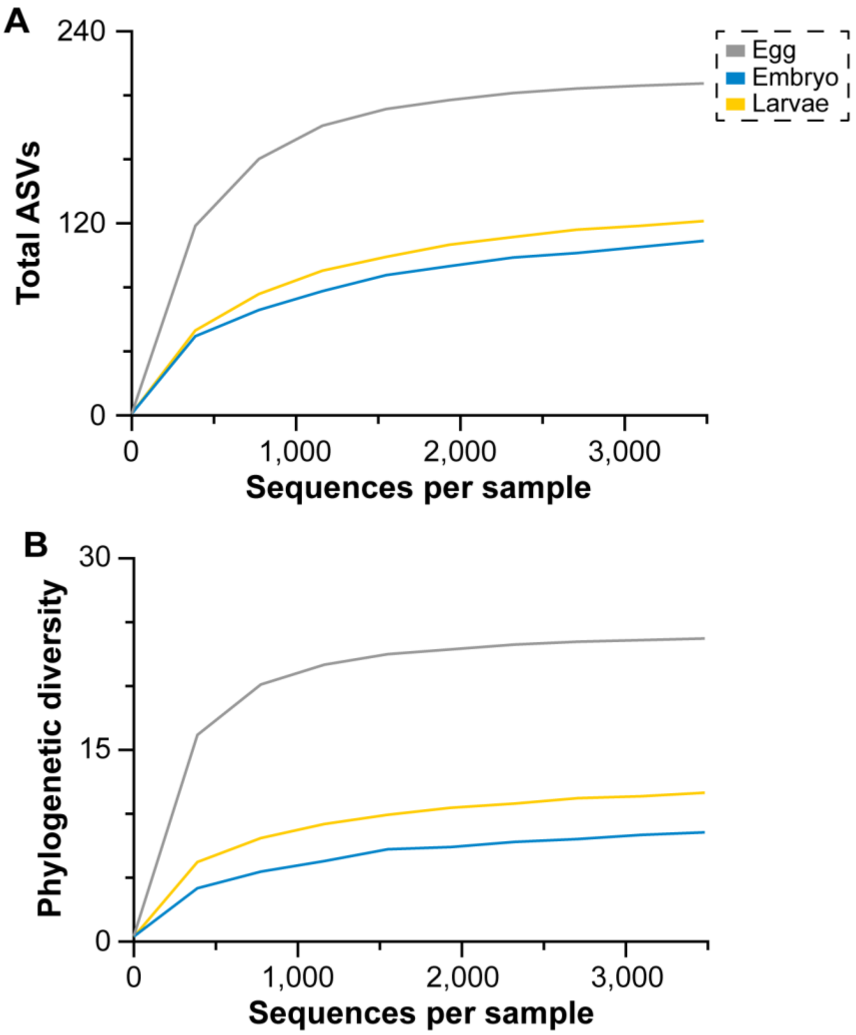


**Fig. S32:** Rarefaction curves for amplicon sequencing. Rarefaction curves for the total ASVs (A) and phylogenetic diversity (B) of the microbial community associated with the developmental stages of the sea urchin *Arbacia lixula*. This was based on a rarefaction depth of 3,482 sequences and was used for all analyses.

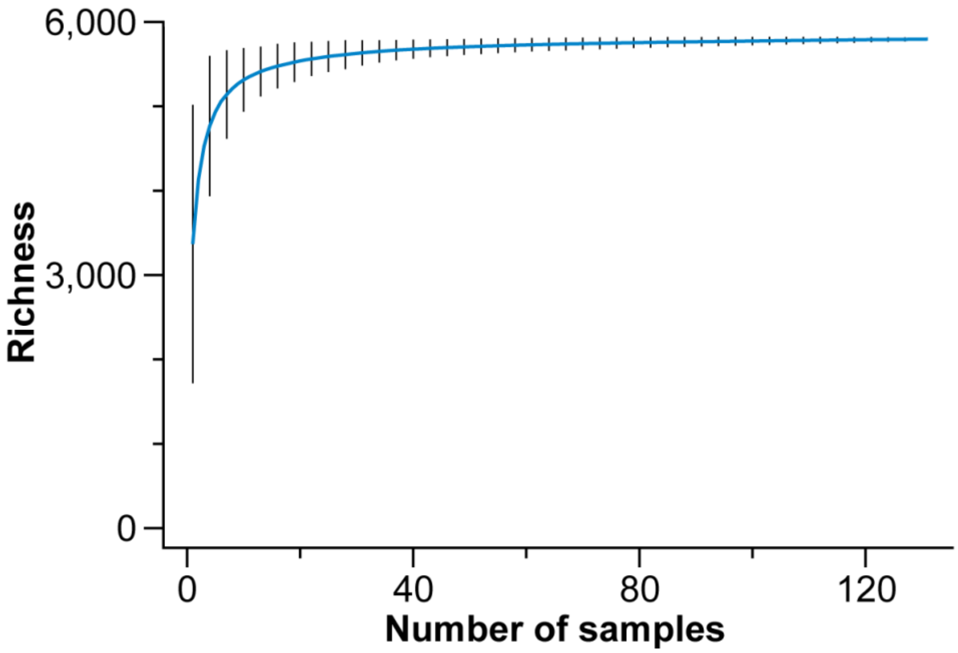


**Fig. S33:** Rarefaction curve for the metabolome. Rarefaction curves for the total metabolites observed based on the samples that were processed for the developmental stages of the sea urchin *Arbacia lixula*.

**Dataset S1A**. Relative abundance of microbial groups in the developmental stages of *Arbacia lixula*.

**Dataset S1B**. Meta-data for meta-analysis of phototrophic eukaryotes associated with sea urchin eggs.

**Dataset S1C**. Relative abundance of Cyanobacteria and photosynthetic eukaryotes that are associated with sea urchin eggs.

**Dataset S1D**. Relative abundance of plastid DNA in sea urchin eggs. This is from the meta-analysis.

**Dataset S1E**. List of plastid ASVs present in sea urchin eggs and their taxonomic assignment. This is from the meta-analysis.

**Dataset S1F**. Beta diversity comparisons between sea urchin eggs and the plastid ASVs that they are provided. This is from the meta-analysis.

**Dataset S1G**. Statistical tables for phylosymbiosis of plastid ASVs. This is from the meta-analysis.

**Dataset S1H**. Ribosomal counts (16S and 18S) from *Arbacia lixula* metagenome of embryos and larvae. Per offspring values were determined by dividing by 200.

**Dataset S1I**. List of metabolites detected in the eggs and larvae of *Arbacia lixula* from light and dark.

**Dataset S1J**. Abundance of each metabolite detected in the eggs and larvae of *Arbacia lixula* from light and dark, as well as controls.

**Dataset S1K**. Summary of statistical tests for sea urchin fitness (*i.e.*, development, morphology, and mortality).

**Dataset S1L**. Summary of statistical tests for metabolomics.

**Dataset S1M**. List of differentially abundance metabolites detected in the larvae of *Arbacia lixula* from light and dark.

**Dataset S1N**. Setting for acquisition during LC-MS/MS metabolomics analysis.
